# Oxidative stress triggers RNAPII arrest through PARylation and DNA damage

**DOI:** 10.1101/2025.11.06.686939

**Authors:** Quentin A. Thomas, Liyang Wu, Emma Lesage, Henriette K. M. Iversen, David López Martínez, Smaragda Kompocholi, Haiyue Liu, Nicolás Nieto Moreno, Lea H. Gregersen

## Abstract

UV or gamma irradiation, as well as certain chemicals, generate DNA damage that disrupts transcription through a variety of well-characterised mechanisms. In contrast, the transcriptional response to oxidative stress remains poorly understood. Here, we describe a rapid and widespread shutdown of transcription following oxidative DNA base damage. By monitoring RNAPII occupancy and elongation dynamics, we demonstrate that oxidative stress temporarily halts RNAPII pause release and arrests the progression of elongation complexes within the gene body. We present evidence that this occurs in a unique and transient manner, characterised by abrupt arrest of elongating RNAPII dead in its tracks, followed by rapid transcriptional recovery as DNA lesions are repaired. We find that the restriction of initiation and early elongation complexes is regulated by PARylation, whereas recovery of RNAPII arrested within the gene body requires DNA repair mediated by the base excision repair (BER) and single-strand break repair (SSBR) pathways.

## Introduction

Our genome is continuously exposed to genotoxic insults causing DNA lesions and genomic instability ^1^. Lesions originating from oxidative damage are among the most frequently observed in the germline ^2^ and accumulation of mutations from oxidative damage has been associated with aging, neurodegenerative diseases, and cancer ^1,3^. Actively transcribed genes are particularly vulnerable to genomic instability, and rapid repair within genes is therefore of utmost importance to prevent deleterious long-term effects ^4,5^. DNA repair and transcription are inevitably coupled as they work on the same template. To maintain fidelity of cellular processes, cells frequently respond to genotoxic stress by silencing transcription, either locally or globally ^4,6–9^. In fact, certain types of DNA lesions can serve as a physical roadblock to transcription, causing a complete halt to RNAPII elongation ^7^. For instance, UV-induced bulky DNA lesions directly impede RNAPII progression and necessitate repair through the transcription-coupled nucleotide excision repair pathway (TC-NER) to facilitate restart of transcription ^9–13^.

In contrast, the direct impact on transcription is less clear in the case of non-bulky DNA lesions and single-stranded breaks (SSBs) frequently induced by oxidative damage. While it is known that exposure to oxidative stress results in a global repression of transcription ^8,14,15^, it remains unknown if and how oxidative DNA lesions impact the transcription machinery *in vivo*. Lesions originating from oxidative stress include non-bulky single-base modification such as 8-oxoguanine (8OG), abasic sites, and SSBs. In fact, SSBs is the most frequent type of DNA lesions, with an estimated occurrence of tens-of-thousands per cell per day ^1,16^. Data generated *in vitro* suggest that RNAPII is able to bypass 8OG DNA lesions ^17–19^, while products originating from further oxidation impede RNAPII progression ^20^. In addition, SSBs generated either as a direct result of oxidative damage, or as BER repair intermediates have been shown to impair RNAPII elongation *in vitro* ^21^. However, transcription assays performed with nuclear extracts from hydrogen peroxide-treated cells remained transcriptionally inactive even on an undamaged DNA template, suggesting that DNA lesions by themselves may not be sufficient to explain the global transcriptional repression observed following oxidative stress ^8^.

Besides physical impairment caused by the DNA lesion itself, the DNA damage response induces a coordinated recruitment of factors involved in DNA repair to the sites of damage, which can elicit either local or global repression of transcription ^22,23^. The recruitment of DNA repair factors can be facilitated either directly by the DNA lesion itself, or through recognition of lesion-stalled RNAPII, such as in TC-NER ^7,9^. Interestingly, rather than being caused by the lesion itself, local transcriptional repression around sites of double-strand breaks (DSBs) is dependent on DNA damage signalling involving ATM, DNAPK, and poly(ADP-ribosyl)ation (PARylation)-mediated regulation of RNAPII close to the site of damage as well as changes in the chromatin environment ^24–28^. PARylation also serves to recruit NELF-E to RNAPII stalled at DSBs ^28^. In addition to DSBs, PARylation is also implicated in the recruitment of DNA repair factors to SSB lesions. Here, PARP1-mediated PARylation stimulates recruitment of X-ray repair cross-complementing group 1 (XRCC1) as well as additional factors involved in SSB repair (SSBR) ^23,29,30^. Interestingly, the lack of XRCC1 leads to a failure of transcription recovery following oxidative stress, suggested to involve PARP1 ‘trapping’ on DNA, which would create a physical block consisting of an unresolved repair intermediate rather than the lesion itself ^15^.

In addition to transcriptional repression induced by DNA damage, it is also becoming evident that cellular stress such as heat shock results in widespread repression of transcription ^31–35^. Multiple mechanisms have been reported to be involved in transcriptional attenuation following heat shock, including promoter-proximal pausing of RNAPII ^33^, NELF condensate formation ^36^, and premature transcript termination ^37^.

Using a variety of genome-wide approaches to track the transcriptional response in a temporally and spatially resolved manner, we find that transcriptional activity in response to oxidative stress is dynamically regulated through early RNAPII pause release, coordinated by PARylation in combination with rapid and reversible arrest of RNAPII in the gene body, controlled at the level of DNA damage repair.

## Results

### Oxidative stress induces a rapid transcriptional shutdown

To investigate the immediate transcriptional response to oxidation-induced stress and DNA damage, we treated cells with a brief, sublethal dose of either H_2_O_2_ or menadione, the latter of which leads to intracellular ROS formation (Fig. 1a, Supplementary Fig. 1a-b). Using metabolic labelling of nascent RNA with 4-thiouridine (4sU) or 5-ethynyluridine (EU) in human cells, we tracked the global levels of RNA synthesis during and upon treatment recovery (Fig. 1b-d and Supplementary Fig. 1c-e). Both H_2_O_2_ and menadione led to a pronounced decrease in overall RNA synthesis across several cell lines (Fig. 1b-d, Supplementary Fig. 1c). In the case of brief H_2_O_2_ treatment, this effect was almost immediate, with a rapid recovery of transcription activity (Fig. 1b-c). In contrast, as expected, menadione induced a somewhat milder and temporally delayed response in terms of repression of nascent transcription levels (Fig. 1b-d). Similarly, lower concentrations of H_2_O_2_ (250 μM) also resulted in a delayed reduction of nascent transcriptional activity (Supplementary Fig. 1c). In contrast, a high H_2_O_2_ dose (10 mM) or menadione (2 mM) resulted in a failure to recover normal transcription levels, impaired cell growth, and apoptosis (Supplementary Fig. 1d-f).

**Fig. 1.**
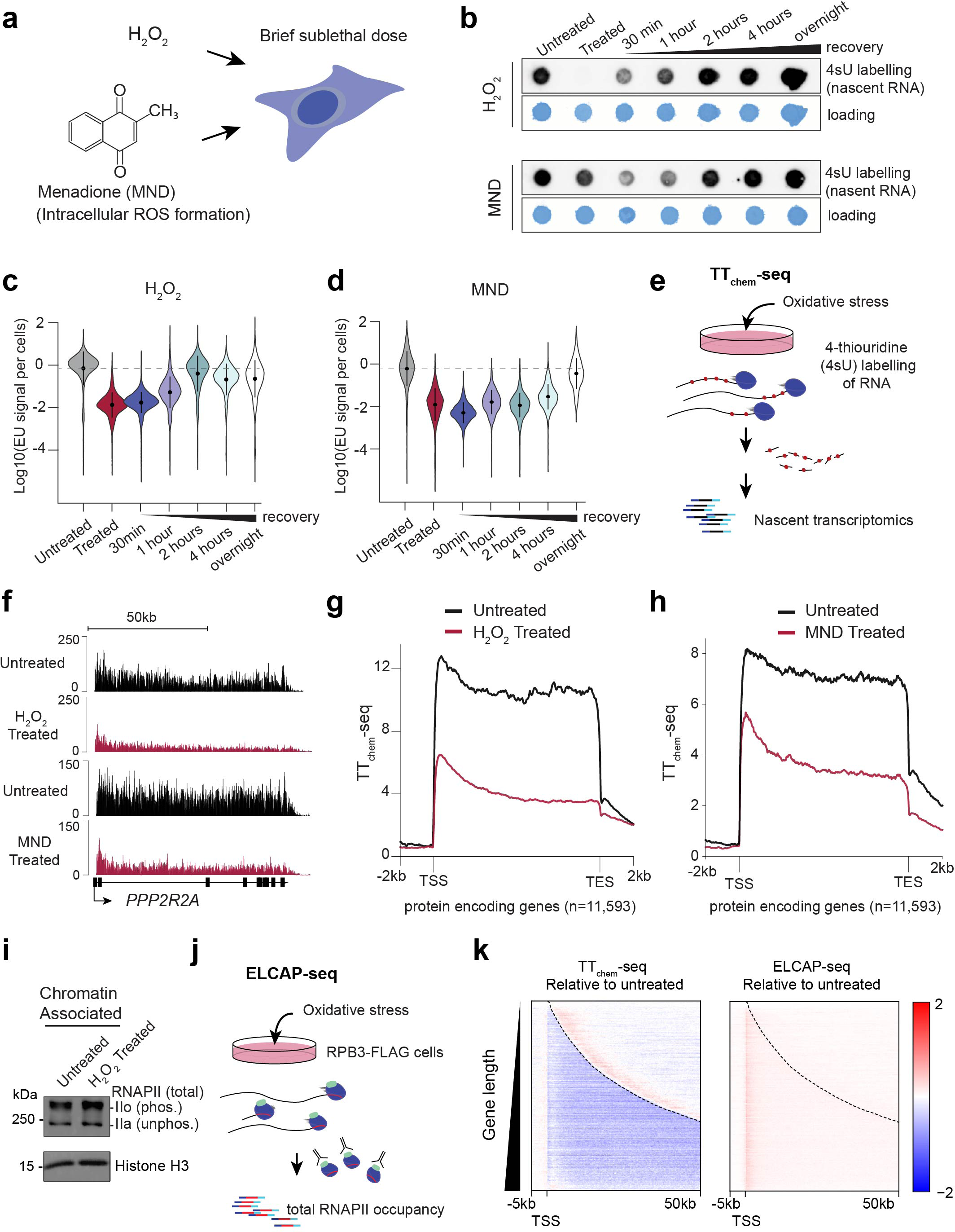
Oxidative stress represses global nascent transcription of RNAPII without release from chromatin. **a** Schematic of oxidative stress inductions using either H_2_O_2_ or Menadione (MND). **b** Nascent transcription levels measured by nascent RNA metabolic pulse labelling (15 min) with 4-thiouridine (4sU). Oxidative stress was induced for 15 min in HEK293 cells with 1 mM H_2_O_2_ or 0.25 mM MND, and cells were allowed to recovery for indicated times. Staining with methylene blue as a loading control. **c-d** Nascent transcription levels quantified by 5-Ethynyluridine (EU) labelling and single cell quantification by immunofluorescence microscopy (EU-assay). Per nuclei EU signal is normalised to DAPI and log10 scaled. Cells treated with H_2_O_2_ (c) or MND (d) and EU-labelled for 30 min. **e** Schematic of TT_chem_-seq profiling nascent transcriptome by 4sU incorporation into nascent RNA, fragmentation and sequencing of purified labelled RNAs. **f** Single-gene view of TT_chem_-seq data between untreated controls and 15 min H_2_O_2_ treated cells and MND treated cells. **g-h** Metagene analysis of TT_chem_-seq data between untreated controls and 15 min H_2_O_2_ treated cells (g) and MND treated cell (h). Lines represent the mean of spike-in normalised read counts between two biological replicates for a set of protein-encoding genes (non-overlapping ± 2 kb, n = 11,593). **i** Western blot of total RNAPII levels from chromatin-enriched cellular fractions. Total RNAPII was detected with an antibody raised against the N-terminal part of RPB1 (D8L4Y). Histone H3 serves as loading control. IIo and IIa denotes hyperphosphorylated and unphosphorylated RPB1 CTD respectively. **j** Schematic of ELCAP-seq experimental design. RNAPII is captured by immunoprecipitation of RPB3-FLAG, short RNA fragments protected by RNAPII are extracted and sequencing to obtain nucleotide resolution maps of RNAPII positions (3’end corresponding to last incorporated nucleotide). **k** Heatmaps of ELCAP-seq signal (left: RPKM) alongside TT_chem_-seq signal (right; read counts normalised to spike-in). The signal is shown as the log2 fold-change between H_2_O_2_ -treated HEK293 cells and untreated cells. Heatmaps centred on transcription start site and sorted by increasing length of gene regions (−5 kb upstream of TSS to 50 kb into the gene body) with dashed lines indicating transcript end sites (TES).

To investigate the effects on transcription genome-wide, we performed TT_chem_-seq ^38^ using yeast spike-ins as an internal normalisation control to accurately quantify changes in nascent transcription in a spatially resolved manner. In agreement with the global quantification from dot blots and EU assays, a dramatic reduction in overall transcription activity throughout the entire gene body of protein-encoding genes was observed (Fig. 1f-h). This was the case both for menadione and for H_2_O_2_ (Fig. 1f-h). In the case of H_2_O_2_, 99.5 % of differentially expressed protein-encoding genes were downregulated (Supplementary Fig. 1g). At lower H_2_O_2_ concentrations, transcriptional repression was strongest at the 30 min recovery timepoint, while a lower menadione concentration led to the strongest repression at 1-hour recovery (Supplementary Fig. 2a-c), which again aligns with the quantifications from dot blots and EU assays (Fig. 1b-c). Classical stress-response genes such as the immediate early genes *FOS* and *EGR1* were also repressed upon H_2_O_2_ and menadione treatment and with kinetics mimicking the global repression (Supplementary Fig. 2d-e), highlighting that even genes later activated as part of various stress-response pathways do not initially escape the transcriptional repression following oxidative stress.

To investigate the possibility that direct inhibition of RNAPII activity might occur because of its oxidation, we assembled *in vitro* transcription elongation complexes with purified RNAPII, pretreated them with increasing concentrations of H_2_O_2_ and performed *in vitro* transcription assays. Whereas the transcriptional activity of RNAPII elongation complexes was reduced at extremely high concentrations of H_2_O_2_ (>50 mM), we failed to observe any impairment of RNAPII activity with lower (<5-10 mM) H_2_O_2_ concentrations, making it unlikely that the lack of transcriptional activity in cells is due to direct RNAPII inactivation (Supplementary Fig. 3a-b).

The results above indicate that transcriptional activity is dramatically reduced throughout the gene body almost immediately after exposure to oxidative stress. The dramatic decrease in nascent transcription within gene bodies can either be due to (1) dissociation of RNAPII elongation complexes from the DNA template, resulting in fewer polymerases in the gene body, or (2) stalling or arrest of RNAPII resulting in disruption of RNA synthesis by those RNAPII elongation complexes. Interestingly, the reduction of transcription did not at first glance appear to be caused by changes in the level of DNA-associated RNAPII as both hyperphosphorylated and unphosphorylated RNAPII levels in chromatin remained unchanged (Fig. 1i).

To more precisely track RNAPII occupancy, we used ELCAP-seq ^39^ which maps RNAPII binding at nucleotide resolution (Fig. 1j). To ensure that all RNAPII populations were effectively captured regardless of their CTD-phosphorylation status, we used cells expressing FLAG-tagged RPB3 (Fig. 1j and Supplementary Fig. 3c). Short RNA fragments protected from nuclease digestion were captured by FLAG-immunoprecipitation of total RNAPII followed by sequencing (Supplementary Fig. 3c-e). Despite the fact that transcription activity was markedly down during the initial response, the levels of RNAPII in the gene body remained largely unchanged (Fig. 1k). The only differences being a slight accumulation of early elongation complexes close to the TSS (Fig. 1k). Intriguingly, this suggests that in response to oxidative stress, elongating RNAPII complexes within the gene body are not dissociated but instead rapidly arrested, in a global manner leading to an immediate transcription impairment.

### Oxidative stress represses RNAPII elongation within the gene body

To confirm that the transcriptional repression observed after oxidative stress is indeed due to reduced transcription in the gene body, we assessed the effect of oxidative stress on gene body-associated RNAPII in the presence or absence of the CDK9 inhibitor 5,6-Dichloro-1-β-D-ribofuranosylbenzimidazole (DRB) (Fig. 2a). In these experiments, RNAPII was first cleared from the gene body by a treatment with DRB for 3.5 hours, which prevents promoter proximal pause release, while allowing all RNAPII to run until the 3’ end of genes and terminate. DRB inhibition is reversible, enabling release of RNAPII complexes into the gene body upon DRB washout. After the 3.5 hours DRB incubation, RNAPII was allowed to enter the gene body by 15 min of unperturbed transcription (RNAPII release phase), after which cells were treated with H_2_O_2_ alone, or in combination with fresh DRB, and incubated for another 15 min (Fig. 2a). This results in the release of RNAPII complexes into the gene body, which are in all cases allowed to run for a total of 30 min (15 min unperturbed + 15 min with indicated treatments). To investigate the effect of oxidation on RNAPII complexes already in the gene body, we focused our analysis on long genes (> 90 kb). In agreement with previous data showing that RNAPII travels approximately 2 kb/min and with a maximum speed around 4 kb/min ^38^, we observed that the RNAPII wavefront progressed to around 60-90 kb into the gene body in cells not treated with H_2_O_2_ (Fig. 2b-e, blue tracks). By contrast, progression of the wave front into the gene body was clearly reduced in response to a brief oxidative stress treatment (Fig. 2b-e, red tracks). Importantly, the wavefront reached a similar point in the gene whether new transcription was inhibited by DRB or not at the time of oxidative stress treatment (Fig. 2b-e; most easily seen by comparing red tracks in d and e). This strongly indicates that H_2_O_2_ treatment also arrested RNAPII molecules that were already in the gene body.

**Fig. 2.**
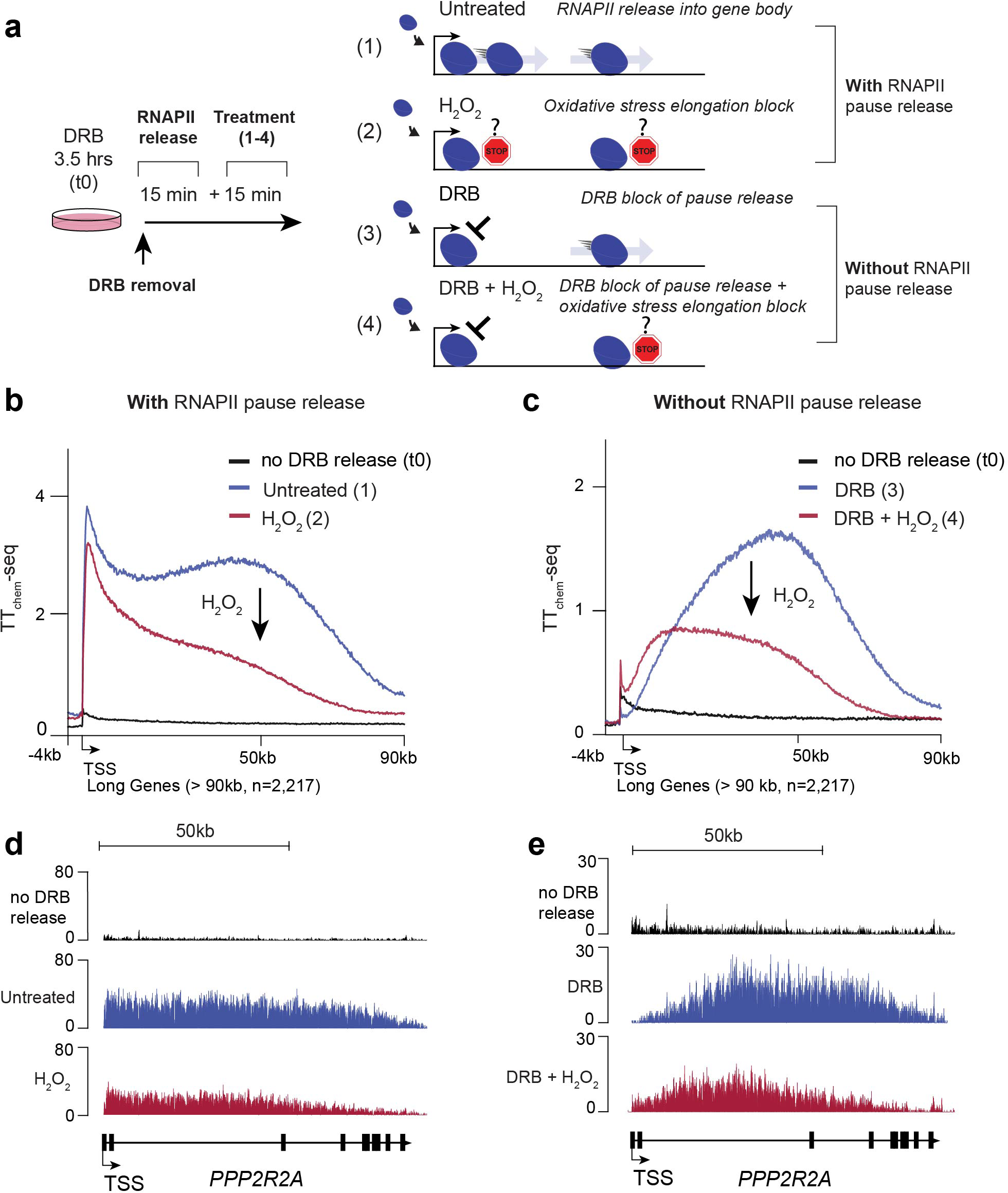
Oxidative stress arrest RNAPII within the gene body. **a** Schematic for RNAPII promoter-proximal release time points with DRB inhibition and release with and without H_2_O_2_ treatment in HEK293 cells. RNAPII complexes are synchronised at promoter-proximal pause site by DRB treatment for 3.5 hours, then released by media washout following either re-inhibition or not and either treated or not with H_2_O_2_. **b** Metagene analysis of TT_chem_-seq data following 3.5 hours of DRB inhibition between no DRB release, 30 min released control and 30 min release with 15 min H_2_O_2_ treatment of HEK293 cells. Data is represented as the average spike-in normalised signal of two biological replicates. The arrow represents the change upon H_2_O_2_ treatment. **c** Metagene analysis of TT_chem_-seq data following 3.5 hours of DRB inhibition with DRB release for 15 min and re-inhibition for 15 min with and without 15min H_2_O_2_ treatment. The arrow represents the change upon H_2_O_2_ treatment. Data is represented as the mean of spike-in normalised TT_chem_-seq read counts over each segmented bins for the selected genes set of protein coding genes and average of two biological replicates. **d-e** Single-gene view of TT_chem_-seq data for the *PPP2R2A* gene for experimental conditions of (b-c) respectively.

### Arrested RNAPII complexes are not subject to ubiquitination and subsequent degradation

A hallmark of RNAPII stalled at bulky UV lesions is polyubiquitination of the largest RNAPII subunit RPB1 and subsequent RNAPII degradation ^10,11^. Unlike the rapid transcriptional repression that occurs following oxidative damage, UV-induced transcriptional repression peaks around 45 min to 3 hrs after the initial exposure (Supplementary Fig. 4a). To examine if RNAPII becomes ubiquitinated following oxidative damage, we tested RPB1 polyubiquitination in a time-course-dependent manner using the DSK2 affinity enrichment strategy for polyubiquitinated proteins ^40^. Although we did observe low levels of RPB1 polyubiquitination, these are much lower than RPB1 polyubiquitination after UV irradiation (Supplementary Fig. 4b). Notably, unlike the case with UV ^10^, we did not observe any decrease in overall levels of hyperphosphorylated transcriptionally engaged RNAPII during either the oxidative stress-induced RNAPII arrest or recovery (Supplementary Fig. 4c).

Upon UV-irradiation, RNAPII stalls at bulky DNA lesions which serve as a roadblock for polymerase elongation ^7^. Accordingly, removal of bulky DNA lesions by the transcription-coupled nucleotide excision repair (TC-NER) pathway is a prerequisite for transcriptional restart following UV ^10,13,41^. Interestingly, TC-NER has also been implicated in lesion repair in response to oxidative stress ^42–45^. To investigate the effects of NER following oxidative stress, we tracked nascent transcription recovery in *XPC* and *CSB* knockout cells, which lack GG-NER and TC-NER, respectively ^7^. Knockout of *XPC* or *CSB* had little or no effect on the transcriptional response (Supplementary Fig. 4d). This is in stark contrast to the transcriptional recovery after UV, which is completely CSB-dependent ^10,12,41^. We also failed to detect recruitment by IP–mass spectrometry of CSB, XPC and other DNA repair factors involved in NER to RNAPII complexes following oxidative stress (Supplementary Table 1). Together with the lack of significant RPB1 ubiquitination observed following oxidative stress (Supplementary Fig. 4b), this indicates that RNAPII arrest following oxidative damage is mechanistically distinct from RNAPII arrest observed as a consequence of UV-irradiation.

### Gene body stalled RNAPII resumes transcription upon recovery from oxidative stress

The data above indicated that in response to oxidative stress, elongating RNAPII complexes are rapidly arrested, dead in their tracks, leading to a global, temporary shutdown of nascent transcription. To further explore the transcription dynamics during the recovery of transcription, we tracked nascent transcription activity over time immediately after oxidation (Fig. 3a). In agreement with the overall transcription levels indicated by dot blots and EU assays, we observed increasing restart of nascent RNAPII transcription 30 min and 1 hour following H_2_O_2_ removal (Fig. 3a-b, Supplementary Fig. 5a). Interestingly, transcription activity was restored through the gene body at both time-points, with higher transcription activity restoration towards the 3’end of long genes at the 1-hour timepoint (Fig. 3a-b, Supplementary Fig. 5b). This again indicates that even RNAPII elongation complexes arrested by oxidation inside the gene body can restart transcription. This point is clearly illustrated by tracking nascent transcription recovery for individual long genes, such as *FARS2* (>570 kb) (Fig. 3b). As stated previously, RNAPII elongation rate measurements indicate that RNAPII travels at approximately 2 kb/min and with a maximum speed around 4 kb/min ^38^. So, if new nascent transcription during the recovery period originated only from newly initiated RNAPII or RNAPII at the promoter-proximal pause site, all transcriptional activity would be restricted to the first 60-120 kb after 30 min recovery and 120-240 kb after 1 hour recovery (Fig. 3b, indicated by arrows). However, recovery of transcription activity was observed throughout the gene already at 30 min and 1 hour recovery (Fig. 3b, indicated by dashed boxes), suggesting that RNAPII complexes indeed resume transcription from within the gene body following oxidation-induced transcriptional arrest.

**Fig. 3.**
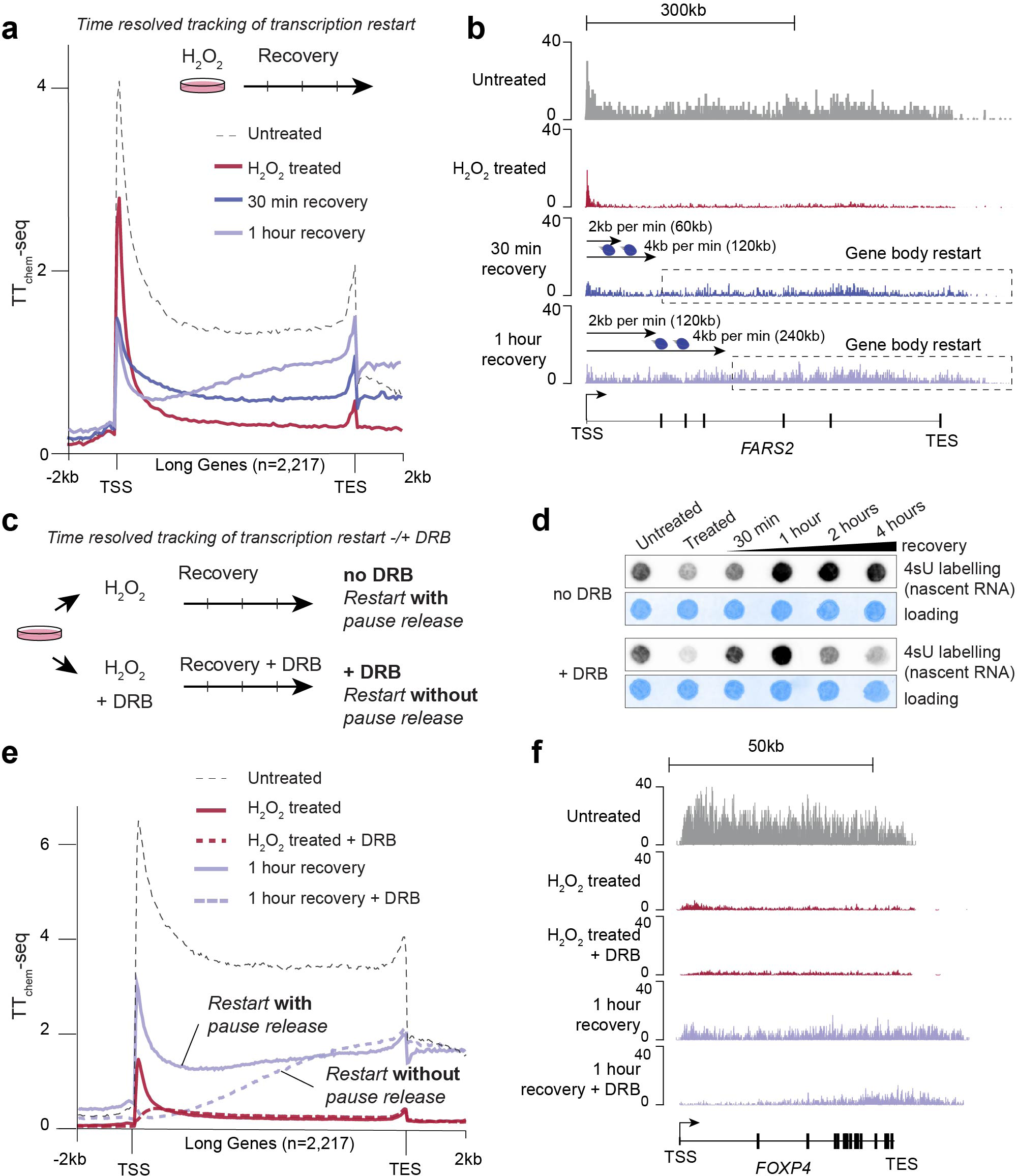
Arrested RNAPII elongation complexes resume transcription from within the gene body during recovery from oxidative stress. **a** Metagene analysis of TT_chem_-seq data for long genes (>90 kb, n=2217) following H_2_O_2_ treatment and indicated recovery time points in HEK293 cells. The line represents the mean of spike-in normalised read counts from three biological replicates. **b** Single-gene view of TT_chem_-seq data for *FARS2* gene (length = 521 kb) for experimental conditions of (a). Arrow represent distance RNAPII would travel from the TSS with an elongation speed of 2 kb/min or 4 kb/min, within the 30 min or 1 hour recovery period, respectively. **c** Schematic of the experimental setup used to study RNAPII restart following oxidative stress with or without DRB inhibition. **d** Nascent transcription levels (4sU dot blot) after 15 min treatment of HEK293 cells with 1 mM H_2_O_2_ with and without DRB. Staining with methylene blue serves as a loading control. **e** Metagene analysis of TT_chem_-seq data for long genes (>90 kb, n=2217) of untreated and 1 hour recovery timepoint following H_2_O_2_ treatment of HEK293 cells, with and without DRB inhibition. Lines is the mean of spike-in normalised read counts between two biological replicates. **f** Single-gene view of TT_chem_-seq data for *FOXP4* gene for experimental conditions of (c).

To further rule out the possibility that newly initiated transcription is responsible for the recovery of transcriptional activity in the gene body, we finally tracked recovery in the absence of new release of RNAPII into the gene body by treating cells with DRB during the recovery period (Fig. 3c). We thus found that overall transcription activity was restored even when DRB was present during recovery from H_2_O_2_ treatment, clearly seen at the 30 min and 1 hour time points (Fig. 3d) before already transcribing polymerases have run off the gene which takes place during longer DRB incubations (Fig. 3d, lower panels). Using TT_chem_-seq, we further confirmed that RNAPII complexes indeed were able to restart transcription from within the gene body in the presence of DRB. Indeed, as expected, transcriptional recovery for one hour after oxidative stress was shifted towards the 3’end and absent in the 5’ end of genes in the presence of DRB (Fig. 3e, dashed purple line; Fig. 3f, lowest panel), and obvious from quantification of nascent transcription levels in the first 25 kb compared to nascent transcription levels in the last 25 kb of genes only (Supplementary Fig. 5b). At the same time, the most prominent change in RNAPII occupancy was a slightly increased pausing index (indicating accumulation of early RNAPII complexes) measured with ELCAP-seq during the recovery phase (Supplementary Fig. 5c). This indicates that the response to oxidative damage is fundamentally different from transcriptional repression induced by bulky DNA lesions which results in RPB1 ubiquitination, subsequent RNAPII degradation and transcription restart as a 5’ to 3’ wave front originating from the TSS by transcription from newly initiated RNAPII complexes ^7,10,46–48^.

To investigate if RNAPII elongation rates change during the recovery phase across genes, we computed the elongation index as the ratio of active elongation (TT_chem_-seq signal) and RNAPII occupancy (ELCAP-seq), which was previously used as a proxy for RNAPII speed ^49,50^. As expected, this indicated a strong decrease in the RNAPII elongation index during the H_2_O_2_ treatment (Supplementary Fig. 5d). As transcription resumes, the RNAPII elongation index in the gene body increased, while the elongation index in the beginning of the gene decreased slightly at the 30 min time point (Supplementary Fig. 5d). At 1 hour recovery this was even more pronounced, with the elongation index almost back to normal levels in the 3’end of the gene body, while the elongation index for early gene body RNAPII complexes decreased even further (Supplementary Fig. 5d). The inverse behaviour of RNAPII elongation indices for early RNAPII complexes and gene body RNAPII indicate that early stalled RNAPII complexes and gene body-arrested RNAPII complexes resume transcription differently during the recovery period (Supplementary Fig. 5c-d). This could either be due to distinct mechanisms controlling release of early RNAPII elongation complexes and gene body-arrested RNAPII, or due to differently configured RNAPII complexes (bound by different RNAPII associated factors) impacting elongation kinetics of RNAPII complexes differently depending on their position at the time of transcription block.

### NELF mediated pausing of RNAPII is altered during the oxidative stress response

To investigate whether RNAPII complexes are remodelled during the global transcriptional repression caused by oxidative stress, we investigated the RNAPII interactome via proteomic analysis of RNAPII immunoprecipitates. We used two approaches (1) capturing total RNAPII complexes using a FLAG-tagged RPB3 cell line that allows us to capture all RNAPII complexes regardless of the phosphorylation status of the CTD, and (2) specifically capturing transcriptionally engaged RNAPII using the monoclonal 4H8 antibody, which recognizes CTD hyperphosphorylated RNAPII (Supplementary Fig. 6a). Notably, using both approaches we observed an increased enrichment of NELF bound RNAPII complexes in H_2_O_2_ treated cells (Fig. 4a-b and Supplementary Fig. 6b, Supplementary Table 1). In addition, we found an increase in 5’ capping components such as CMTR1 and RNGTT associated RNAPII complexes upon treatment with H_2_O_2_ (Fig. 4a-b). This is in line with data suggesting that recruitment of capping factors is dependent on NELF ^51^ and suggests increased early RNAPII pausing. We confirmed the increased interaction between RNAPII and the NELF complex by RNAPII IP-western analysis (Fig. 4c). Interestingly, while NELF recruitment to RNAPII was initially increased, it decreased during the recovery period and after 1 hour and 2 hours of recovery, NELF was less associated with RNAPII than even the untreated conditions (Fig. 4c). This trend was also observed in RNAPII IP proteomics both at 1 hour and 2 hours recovery (Supplementary Fig. 6b).

**Fig. 4.**
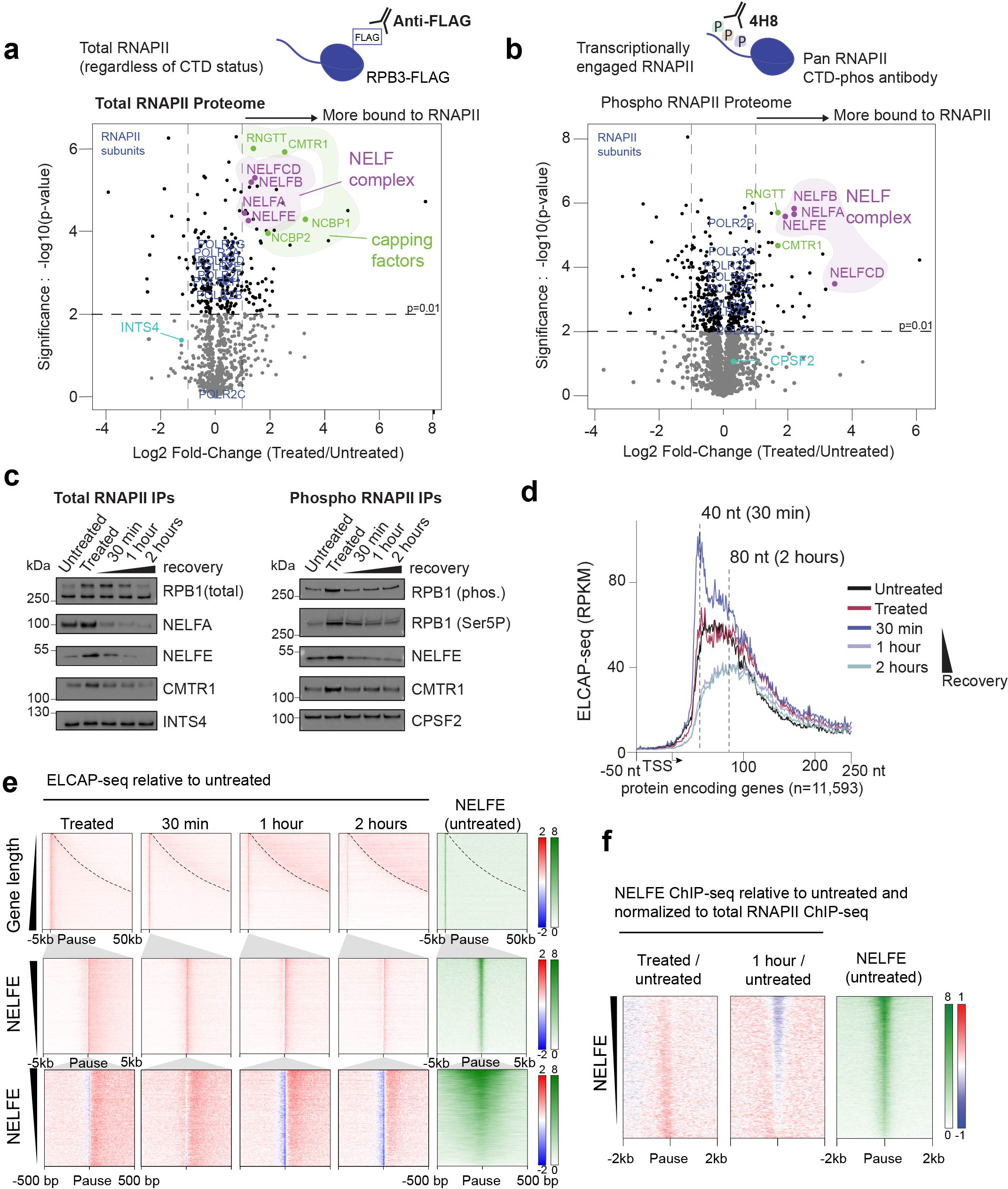
NELF mediated RNAPII pausing is altered during the oxidative stress response. **a** Total RNAPII (FLAG IP in HEK293 RPB3-FLAG cells) interactome after 15 min of 1 mM H_2_O_2_ treatment. A log2 fold-change comparison of H_2_O_2_-treated with untreated control cells is shown relative to significance as -log2 of p-value (triplicate injections). Are highlighted: RNAPII, NELF and the capping complex subunits and INTS4 (IP-WB loading control in (c)). **b** Phosphorylated RPB1 (4H8 IP, HEK293 cells) interactome after 15 min of 1 mM H_2_O_2_ treatment. A log2 fold-change comparison between H_2_O_2_ treated to untreated control cells is shown relative to significance as -log2 of p-value. Are highlighted: RNAPII subunits, NELF subunits, members of the capping complex and CPSF2 (IP-WB loading control) in (c) are coloured and labelled. **c** Western blot analysis of FLAG IP (left) and 4H8 IP (right) before and after H_2_O_2_ treatment. Total RNAPII probed with D8L4Y antibody. Phosphorylated RNAPII CTD with 4H8. Serine 5 phosphorylated RNAPII CTD with 3E8. **d** Metagene analysis of ELCAP-seq, RPKM signal centred on the TSS for non-overlapping coding genes (n=11,593) in untreated control, H_2_O_2_ treated, and recovery timepoints (30 min, 1 hour and 2 hours). The lines are the average signal of two biological replicates. Doted vertical lines are the highest RNAPII occupancy distance from TSS upon 30 min recovery and 2 hours recovery. **e** Heatmap analysis of RPKM ELCAP-seq in RPB3-3xFLAG cells treated with H_2_O_2_ as in (d) and NELFE untreated ChIP-seq signal in HEK293. The ELCAP-seq signal is shown as a log2 fold-change relative to untreated. NELFE ChIP-seq signal is normalised to IgG input. Top panel shows gene regions (pause −5 kb to pause +50 kb) sorted by increasing gene length (pause site genes n= 7,595), with dashed lines indicating TES. Middle panel shows promoter regions (pause site ± 5 kb) sorted by decreasing NELFE ChIP-seq signal. Bottom panel shows promoter regions (pause site ± 0.5 kb) sorted by decreasing NELFE ChIP-seq signal. **f** Heatmap of NELFE ChIP-seq log2 fold-change between H_2_O_2_ treated and untreated control and between 1 hour recovery and untreated control, both normalised by ratio to the log2 fold-change of respective treatment changes in total RPB1 (D8L4Y) ChIP-seq in HEK293 cells. Gene regions (± 5 kb pause site genes n = 7,595) are sorted by decreasing NELFE ChIP-seq signal in untreated condition.

To address the role of NELF-mediated RNAPII regulation during the oxidative stress response, we carried out a detailed analysis of RNAPII occupancy. ELCAP-seq provides nucleotide resolution of RNAPII binding and is therefore ideally suited to investigate RNAPII promoter-proximal pausing. This uncovered an increase in promoter-proximal pausing at 30 min after oxidative stress (Fig. 4d and Supplementary Fig. 6c). Interestingly, pausing after 30 min recovery was primarily increased at the canonical pause site (20-50 bp from the TSS), also known as the 1^st^ pause site of NELF-mediated pausing ^51,52^. However, at later time points (1 hour and 2 hours recovery), RNAPII occupancy at the canonical pause site decreased, with the RNAPII occupancy shifting downstream to the ‘2^nd^ pause site’ coinciding with the +1 nucleosome ^51^ (Fig. 4d-f). Tellingly, a similar shift from 1^st^ to 2^nd^ pause site has previously been described to occur after loss of NELF ^51,52^. To test if the increased association between NELF and RNAPII correlates with the changes in promoter-proximal pausing, we plotted the relative NELF binding (ascertained by NELFE ChIP-seq) after normalisation to RNAPII binding levels (spike-in normalised RNAPII ChIP-seq) (Fig. 4f). During the initial repression after oxidative stress, more NELF accumulated at the promoter-proximal pause site relative to RNAPII, indicating active NELF recruitment, or impaired NELF release, at paused RNAPII complexes (Fig. 4f). However, during recovery, less NELF was bound to promoter-proximal RNAPII, in agreement with our IP results and the observed shift in RNAPII pausing from the 1^st^ pause site to the 2^nd^ pause site (Fig. 4c-f).

We wondered if the increased NELF binding to RNAPII during the initial phase of the oxidative stress response serves to restrict paused RNAPII elongation complexes from entering productive elongation. To test this, we generated NELFC degron cell lines enabling us to track nascent transcription in response to oxidative stress in cells lacking the NELF complex (Supplementary Fig. 6d). Treatment with dTAG led to depletion of NELFC within 1 hour (Supplementary Fig. 6e). As previously reported ^51^, NELFC depletion also resulted in degradation of additional NELF subunits, such as NELFA (Supplementary Fig. 6e). Intriguingly, however, the overall levels of nascent transcription either during the immediate response to H_2_O_2_ or in the recovery period were unaffected by loss of NELF (Supplementary Fig. 6f). As NELF condensation formation has been suggested to drive transcriptional repression during the heat shock response ^36^, we checked nuclear localisation of NELF during the oxidative stress response. As expected from previous data by others, changes in the nuclear location of NELF condensate formation was observed following heat shock, but no indications of condensate formation following oxidative stress were observed (Supplementary Fig. 6g). Thus, the NELF depletion experiments indicate that NELF-mediated regulation is not responsible, nor required, for the widespread transcriptional repression observed upon oxidative stress. Instead, we observe that the *position* of early RNAPII pausing is affected during the oxidative stress response in a NELF-dependent manner; first accumulating at the 1^st^ pause site (increased NELF binding) followed by a shift towards the more downstream +1 nucleosome pause site (decreased NELF binding).

### PARylation governs early transcriptional recovery

Another means of promoting RNAPII stalling in response to DNA damage is PARylation catalysed by poly(ADP-ribose) polymerase 1 (PARP1) ^14^. PARP1 activation leads to PARylation of PARP1 itself (auto-PARylation) and other proteins which serve as scaffolds for recruitment of DNA repair factors at the site of DNA lesions ^23,53^. To test if PARylation might be relevant for the transcriptional response to oxidative stress, we first measured the level of ADP-ribose on proteins in a time-dependent manner. In agreement with previous data ^14^, we observed a strong and rapid increase in PARylation during the initial oxidative stress response (Fig. 5a, left). The highest levels were observed during the H_2_O_2_ treatment itself, coinciding with the time transcriptional arrest (Fig. 5a, left). During the recovery phase PARylation levels rapidly decreased and almost reached baseline levels at the 2 hours recovery timepoint (Fig. 5a, left). PARylation can be modulated by treatment with either PARP inhibitor (PARPi) or poly(ADP-ribose) glycohydrolase inhibitor (PARGi) ^54^. As expected, treatment of cells with PARPi prior to oxidative stress abolished PARylation (Fig. 5a, right) while pre-treatment with a PARG inhibitor (PARGi) resulted in sustained PARylation (Fig. 5b). To test if PARylation might play a functional role during the oxidative stress response we first tested the overall levels of nascent transcription in cells treated with either PARPi or PARGi prior to the induction of oxidative stress. Strikingly, PARPi led to faster recovery of overall transcription levels, while sustained PARylation following treatment with PARGi resulted in reduced transcriptional recovery (Fig. 5c and Supplementary Fig. 7a-b). To address how PARylation affects nascent RNAPII transcription in a genome-wide manner, we performed nascent transcriptome profiling by TT_chem_-seq. Confirming the dot blot results, we first observed higher levels of overall 4sU-labelled RNA at the 15 min timepoint in PARPi treated cells (Fig. 5d). Somewhat surprisingly, the TT_chem_-seq data revealed that PARPi led to increased release of early RNAPII elongation complexes into the gene body at the 15 min recovery timepoint (Fig. 5e-f). The same trend of increased nascent transcription levels primarily within the first 40 kb upon PARPi pre-treatment was also clearly observed at a single gene level (Fig. 5g) and across genes regardless of gene length (Fig. 5h). With an RNAPII elongation rate of 2-4 kb/min, RNAPII complexes originating from newly initiated RNAPII complexes or released from the promoter proximal site would be expected to travel 30-60 kb into the gene body at the 15 min timepoint, which corresponds nicely to the region in which the primary increase in RNAPII transcriptional activity upon PARPi treatment was observed (Fig. 5e-h). This suggests that PARylation after oxidative stress inhibits either RNAPII initiation or RNAPII promoter-proximal pausing, or both. Strikingly, the spatial effect of PARPi treatment on nascent transcription across the gene was sustained even at the 1-hour recovery timepoint, despite seeing similar overall levels of nascent transcription between PARPi and control-treated cells at this timepoint (compare Fig. 5c and 5h), again indicating that the effect of PARylation is primarily restricted to early elongation.

**Fig. 5.**
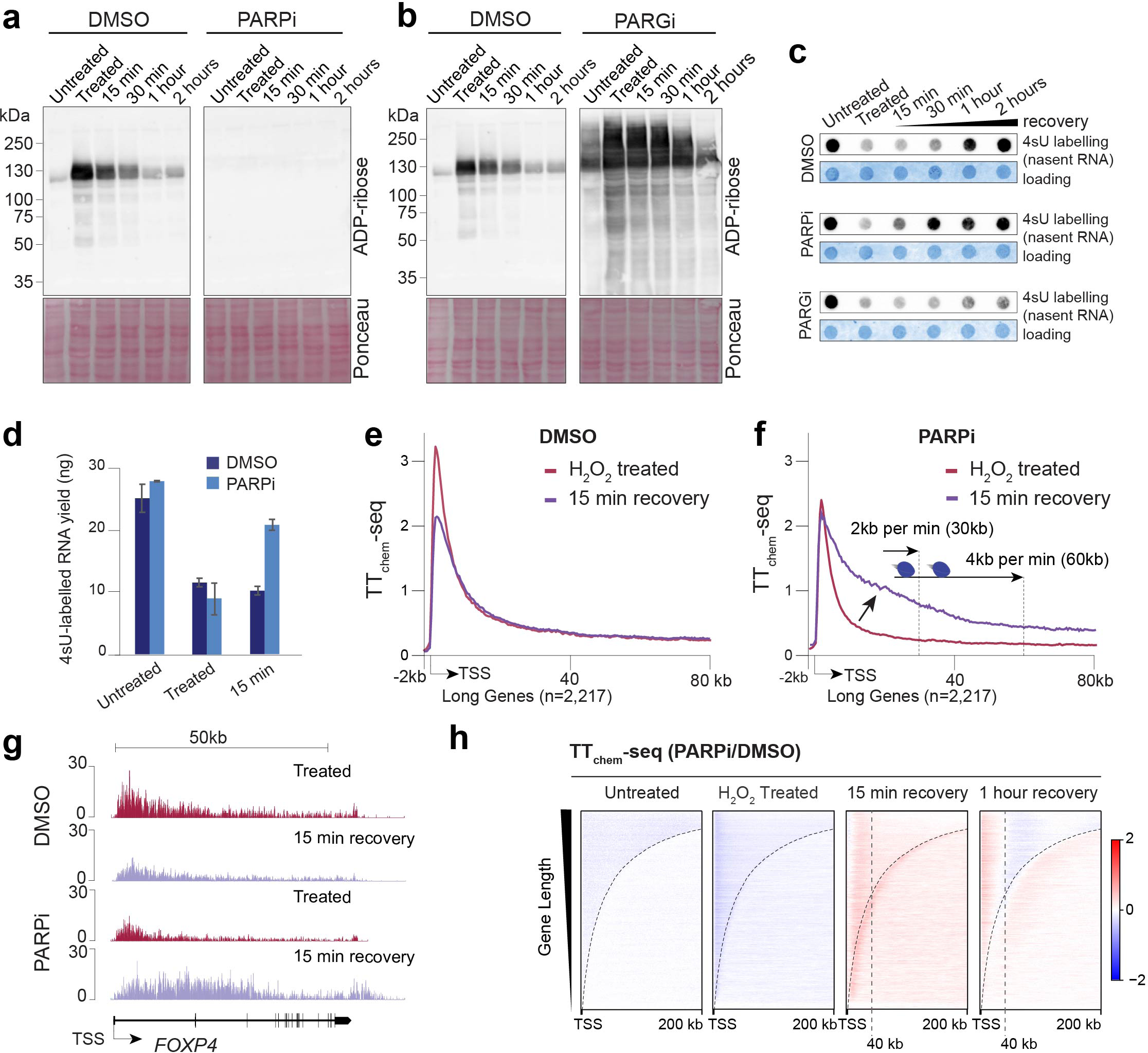
PARylation restricts release of early RNAPII elongation complexes during transcription recovery. **a-b** Western blot analysis of poly-ADP-ribose levels in HEK293 cells with and without 1 hour pre-treatment with PARP inhibitor (Olaparib) (a) or cells pre-treated for 1 hour with PARPG inhibitor (PDD 00017273) (b) and then subsequently treated with 1 mM H_2_O_2_ for 15 min and indicated times following media replenishment. Ponceau S staining serves as total protein loading control. **c** Nascent transcription levels (4sU dot blot) treated as in (a) with and without PARPi pre-treatment or PARGi. Staining with methylene blue serves as a loading control. **d** 4sU labelled RNA yields measured after streptavidin pull-down preformed in duplicates of HEK293 cells treated as in (a), with or without 1 hour PARPi pre-treatment. **e-f** Metagene analysis of TT_chem_-seq for long genes (non-overlapping >90 kb, n = 2,217) of DMSO treated HEK293 cells (e) or PARPi treated HEK293 cells (f) treated with 1 mM H_2_O_2_ for 15 min with and without 15 min recovery. Lines is the mean of spike-in normalised read counts between two biological replicates. **g** Single-gene view of TT_chem_-seq data for the *FOXP4* gene for experimental conditions of (E-F). **h** Heatmaps of TT_chem_-seq signal (normalised to spike-in). The signal is shown as the log2 fold-change between PARPi treated HEK293 cells and DMSO treated cells as control. Heatmaps are centred on transcription start site and are sorted by decreasing length of genes regions (TSS to +200 kb) with dashed lines indicating TES.

This differential effect indicates that the transient arrest of RNAPII further into the gene body relies on other mechanisms or additional factors beyond PARylation signalling. Intriguingly, NuMA, a nuclear protein associated with the mitotic spindle, binds RNAPII in a PARP1-dependent manner with repair proteins TDP1 and XRCC1 and has been suggested to help promote oxidative break repair ^55^. In apparent agreement with the notion that PARylation promotes NuMA binding ^55^, we indeed observed increased association of NuMA with RNAPII immediately after oxidative stress (Supplementary Table 1). Importantly, however, NuMA was not detected in RNAPII elongation complexes captured by IP of hyperphosphorylated RNAPII, but only in IPs capturing total RNAPII complexes (Supplementary Table 1). This is in agreement with the idea that NuMA associates primarily with initiating RNAPII, and in line with the findings above that PARylation primarily affects this RNAPII population.

### Lack of SSBR prevents transcriptional recovery

A key function of PARP activation in response to oxidatively induced DNA damage is to facilitate recruitment of the molecular scaffold protein XRCC1 to the single-stranded DNA breaks (SSB) generated either as a direct consequence of oxidative damage or by incision by DNA glycosylases such as OGG1, promoting efficient base excision repair (BER) ^29,30^. To investigate the role of XRCC1 in the transcriptional response to oxidative stress, we generated *XRCC1* KOs (Supplementary Fig. 7c). In agreement with previous reports ^15^, *XRCC1* KOs showed a clear defect in recovery of nascent transcription following oxidative stress (Fig. 6a-b). This effect was observed even 2 and 4 hours after treatment, suggesting that XRCC1 is critical for transcriptional recovery following oxidative stress (Fig. 6a-b). Using nascent transcriptomics, we found that lack of XRCC1 led to an almost complete failure of RNAPII transcription restart throughout the gene body (Fig. 6c). Compared to WT cells, *XRCC1* KOs first and foremost displayed much less transcription recovery towards the end of long genes at the 1-hour recovery time point (Fig. 6c), indicative of failure to restart gene body-arrested RNAPII complexes. Previous reports have indicated that XRCC1 is dependent on PARylation for its rapid recruitment onto oxidatively induced SSB ^30^. Noticeably, pre-treatment of *XRCC1* KOs with PARPi rescued the delay in overall nascent transcription levels observed in the *XRCC1* KOs (Fig. 6d). This suggests that PARylation serves a role in the transcriptional response beyond simply facilitating XRCC1 recruitment to sites of DNA damage.

**Fig. 6.**
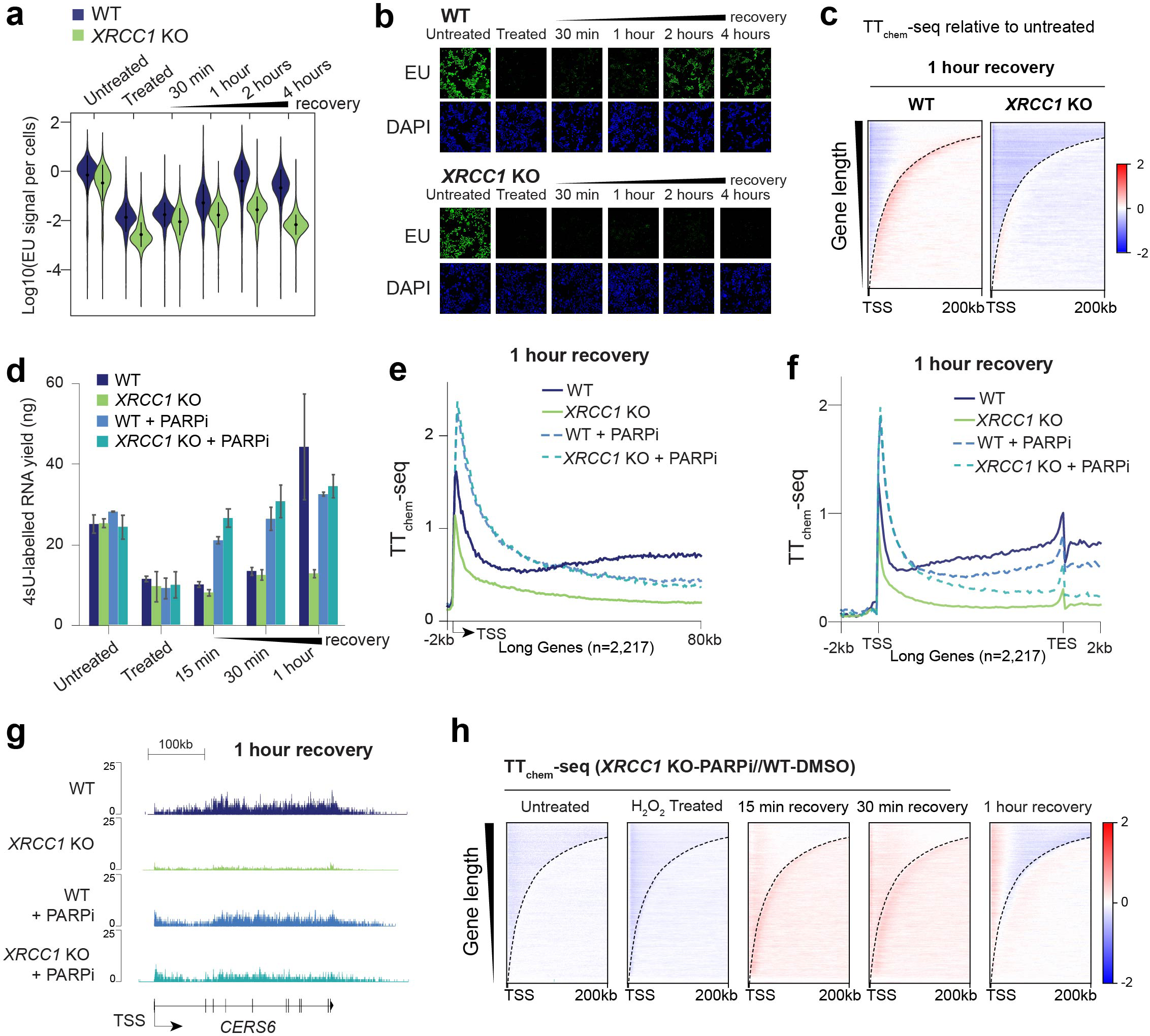
Lack of XRCC1 blocks gene body recovery of RNAPII elongation complexes during transcription recovery. **a** Nascent transcription levels quantified by 5-Ethynyluridine (EU) labelling and single cell quantification by immunofluorescence microscopy (EU-assay). Per cell EU signal is normalised to DAPI and log10 scaled. HEK293 WT and *XRCC1* knockouts (KO) cells were treated with 1 mM H_2_O_2_ and EU-labelled for 30 min. **b** Representative images of global nascent transcription (EU immunofluorescence) from experiments in (a) of WT HEK293 and *XRCC1* KO cells following oxidative stress conditions. All image intensities are displayed with same exposure as the untreated control. **c** Heatmaps of TT_chem_-seq signal (normalised to spike-in) in HEK293 WT cells and *XRCC1* KO cells. The signal is shown as the log2 fold-change between 1 hour recovery relative to untreated samples for the respective cell lines. Heatmaps are centred on TSS and are sorted by decreasing length of gene regions (TSS to +200 kb) with dashed lines indicating TES. **d** 4sU labelled RNA yield measured after streptavidin pull-down in HEK293 cells (WT), *XRCC1* KO (average for two independently generated KO cell lines), HEK293 treated with PARPi (WT + PARPi) and *XRCC1*-KO treated with PARPi (*XRCC1* KO + PARPi). All samples were performed in duplicates. **e-f** Metagene analysis of TT_chem_-seq data for 1 hour recovery for long protein coding genes (>90 kb, n=2,217). Metagene in (e) is centred on TSS (-2 kb to TSS to +80 kb) while metagene in (f) is scaled on transcription start site and transcript end site (-2 kb to TSS to TES +2 kb). Lines is the mean of spike-in normalised read counts between two biological replicates. **g** Single-gene view of TT_chem_-seq data for the *CERS6* gene for experimental conditions of (e-f). **h** Heatmaps of TT_chem_-seq signal (normalised to spike-in). The signal is shown as the log2 fold-change between WT-DMSO control and PARPi treated *XRCC1*-KO cells. Heatmaps are centred at the transcription start site (TSS + 200 kb) and sorted based on gene lengths.

To address the transcriptional interplay between XRCC1 and active PARylation genome-wide, we performed nascent transcriptome profiling by TT_chem_-seq of *XRCC1* KOs, with and without PARPi pre-treatment (Fig. 6e-h). Interestingly, when focusing on elongation close to the TSS, PARPi completely reversed the effect of *XRCC1* KOs at 1 hr recovery (Fig. 6e). Thus, PARPi treatment both in WT and *XRCC1* KOs led to enhanced transcription activity early in the gene body, in accordance with the previous results showing PARylation-dependent release of early RNAPII complexes. However, looking at metagene plots for long genes, it was obvious that PARPi treatment could not fully reverse the transcription recovery towards the 3’end of long genes even at the 1-hour recovery timepointc. 6f). Tellingly, PARPi treatment in *XRCC1* KO cells did not fully rescue the lack of nascent transcription recovery towards the end of genes to the same levels as PARPi treated cells, indicating a PARylation-independent effect of XRCC1 on gene body recovery (Fig. 6f). This is also evident from single gene examples, where transcription recovery in the beginning of *CERS6* is similar for PARPi treatment in WT and *XRCC1* KOs, while 3’end recovery is not fully rescued by PARPi treatment in *XRCC1* KOs (Fig. 6g). Looking specifically at the PARPi effect in *XRCC1* KOs across genes with ranked according to their lengths, we again see an effect of PARPi restricted primarily to the initial 40 kb region (Fig. 6h). This suggests that while PARPi promoted release of early elongation complexes can partially ‘override’ the lack of transcription recovery in *XRCC1* KOs through release of early elongation complexes, PARPi treatment cannot fully overcome transcriptional defects further into the gene body (Fig. 6h). To investigate if the effect of XRCC1 might be due to PARP ‘trapping’ on DNA as previously reported ^56^, we checked the levels of chromatin-associated PARP1 protein in a time-dependent manner. However, we found that PARPi did not led to increased chromatin levels of PARP1, either in WT cells or in *XRCC1* KOs (Supplementary Fig. 7d), arguing that PARP trapping is unlikely to be the cause of the transcriptional phenotypes observed upon PARPi treatment in our experiment.

### Lack of OGG1 activity slows transcriptional restart

One of the most frequent oxidative damage-induced DNA lesions is the non-bulky 8-oxoguanine (8OG), which is primarily recognised by the base excision repair (BER) pathway. Although RNAPII has been shown to bypass 8OG lesions present in naked DNA templates ^17–19^, it is unknown how 8OG might impact the transcription machinery *in vivo*. To investigate the role of BER and whether BER is required for transcription recovery we chemically inhibited the 8OG DNA glycosylase-1 (OGG1). Interestingly, OGG1 inhibition (OGG1i) also led to a clear delay in the recovery of transcription levels (Fig. 7a-b), indicating that recovery of nascent transcription is dependent on the BER pathway. This was again further confirmed by nascent transcriptome profiling by TT_chem_-seq, which showed that transcriptional restart within the gene body at 1 hour recovery was almost absent (Fig. 7c-f). Intriguingly, although treatment with OGG1i delayed restart of RNAPII, the level of transcriptional activity at later time points recovered close to levels of untreated cells (Fig. 7a-b), suggesting that rather than preventing transcriptional restart completely, BER defects such as those inflicted by lack of OGG1 activity delays transcriptional restart. This contrasts with the situation in *XRCC1* KOs where transcription activity remained impaired even at 4 hours recovery (Fig. 6a-b). To investigate how BER defects influence the PARylation-driven release of early RNAPII complexes, we combined OGG1i with PARPi pre-treatment. Firstly, we wanted to see if PARPi treatment could override the dramatic sustained OGG1i nascent transcription repression. Indeed, similarly to the situation for XRCC1, PARPi pre-treatment resulted in recovery of overall nascent transcription levels starting already at the 15 min recovery time point (Fig. 7d) which was maintained throughout the recovery period (Fig. 7d and Supplementary Fig. 7e). To determine if the PARPi effect was restricted to release of early RNAPII elongation complexes, we performed TT_chem_-seq and looked specifically at the nascent transcription recovery for long genes at the 1-hour timepoint. Like the situation for XRCC1, the added transcriptional activity brought about by PARPi pre-treatment in OGG1i treated cells was primarily restricted to early elongation complexes (Fig. 7e). However, when focussing also on the 3’end of long genes, PARPi pre-treatment was able to overcome the OGG1i elongation block to a greater extent than in *XRCC1* KOs (Fig. 7f). Nevertheless, gene body transcription levels in the 3’end of long genes were still greatly reduced compared to WT conditions (Fig. 7e). Something that is also very evident from an individual gene example of the long gene *PAM* (Fig. 7g). The spatially restricted effect of PARPi ‘forced’ early RNAPII release was further confirmed by heatmaps looking across both long and short genes (Fig. 7h). Thus, again PARPi treatment leads to release of early RNAPII elongation complexes, an effect that can partially override the elongation block observed in cells defective for OGG1-mediated repair (Fig. 7e-h, Supplementary Fig. 7e). This again suggests that while PARylation restricts early elongation complexes, DNA repair is crucial for resumption of RNAPII transcription within the gene body. An effect which becomes particularly evident when looking at the 3’end of long genes (Fig. 7f-h). Together, these data thus indicate that, after oxidative damage, PARylation is important to restrict release of RNAPII from promoter-proximal areas, while DNA repair is required to remove DNA lesions in the path of RNAPII.

**Fig. 7.**
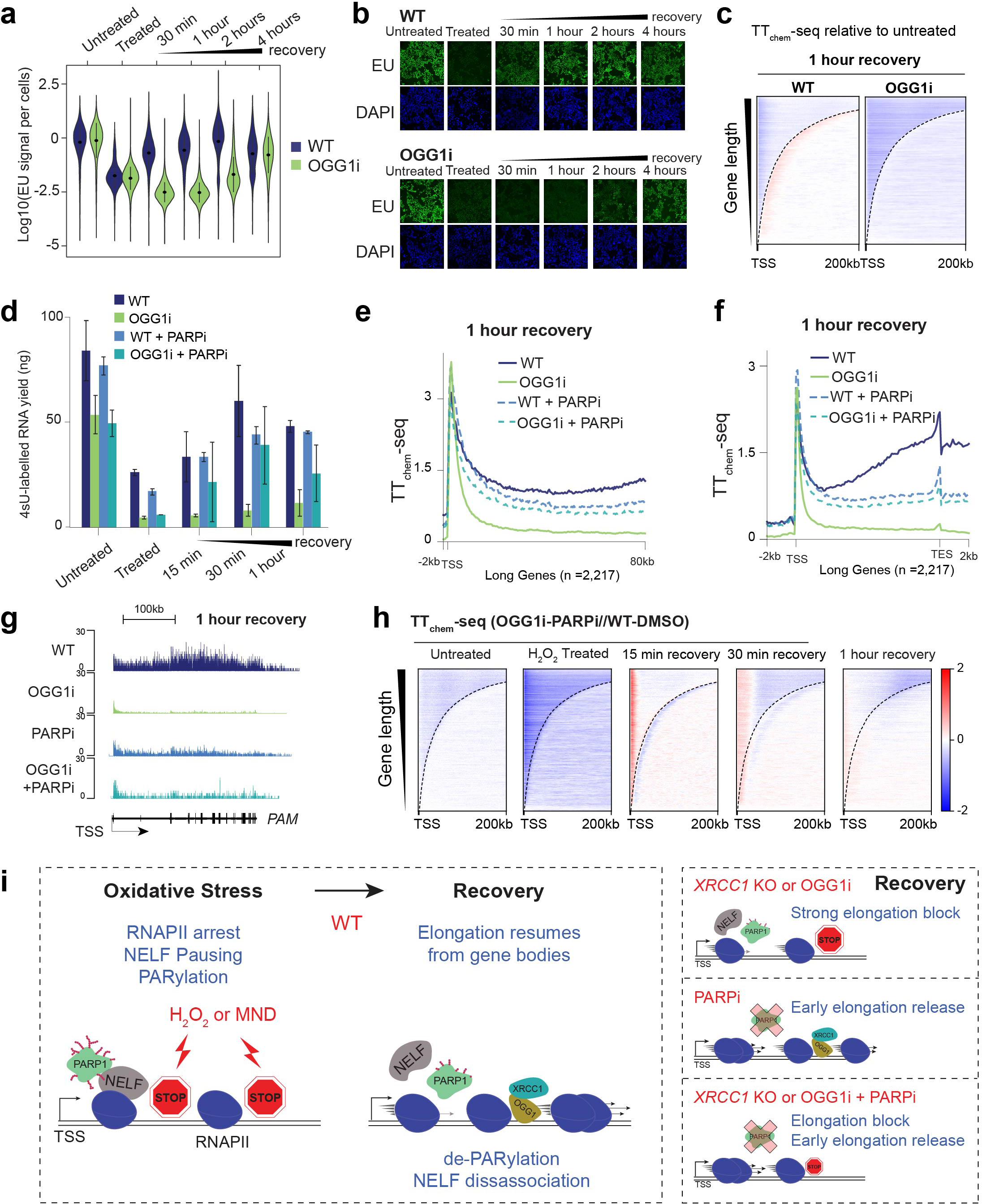
Lack of OGG1 activity blocks gene body recovery of RNAPII elongation complexes during transcription recovery. **a** Nascent transcription levels quantified by 5-Ethynyluridine (EU) labelling and single cell quantification by immunofluorescence microscopy (EU-assay). Per cell EU signal is normalised to DAPI and log10 scaled. HEK293 WT and OGG1i treated HEK293 cells were treated with 1 mM H_2_O_2_ and EU-labelled for 30 min. **b** Representative images of global nascent transcription (EU immunofluorescence) from experiments in (a) of WT HEK293 and OGG1i treated HEK293 cells following oxidative stress conditions. All image intensities are displayed with same exposure as the untreated control. **c** Heatmaps of TT_chem_-seq signal (normalised to spike-in) in HEK293 WT cells and OGG1i treated HEK293 cells. The signal is shown as the log2 fold-change between 1 hour recovery relative to untreated samples for the respective cell lines. Heatmaps are centred on TSS and are sorted by decreasing length of gene regions (TSS to +200 kb) with dashed lines indicating TES. **d** 4sU labelled RNA yield measured after streptavidin pull-down in HEK293 cells (WT), HEK293 treated with OGG1i, HEK293 treated with PARPi (WT + PARPi) and OGG1i treated HEK293 with PARPi (OGG1i + PARPi). All samples were performed in duplicates. **e-f** Metagene analysis of TT_chem_-seq signal for 1 hour recovery for long protein coding genes (>90 kb, n=2,217). Metagene in (e) is centred on TSS (-2 kb to TSS to +80 kb) while metagene in (f) is scaled on transcription start site and transcript end site (-2 kb to TSS to TES +2 kb). Lines is the mean of spike-in normalised read counts between two biological replicates. **g** Single-gene view of TT_chem_-seq data for the *PAM* gene for experimental conditions of (e-f). **h** Heatmaps of TT_chem_-seq signal (normalised to spike-in). The signal is shown as the log2 fold-change between WT-DMSO control and PARPi treated OGG1i treated HEK293 cells. Heatmaps are centred at the transcription start site (TSS + 200 kb) and sorted based on gene lengths. **i** Model of the transcriptional response to oxidative damage. In WT cells (left) oxidative damage induced by hydrogen peroxide or menadione results in rapid RNAPII arrest, NELF-mediated promoter proximal pausing and PARylation. As transcription recovers RNAPII elongation resumes from within the gene body. NELF is disassociated from early RNAPII elongation complexes and released into the gene body due to de-PARylation. In *XRCC1* KOs or upon OGGi RNAPII elongation block is maintained (right top panel), PARPi treatment results in forced release of early RNAPII elongation complexes (right middle panel), whereas PARPi in *XRCC1* KOs or OGG1i leads to forced release of early RNAPII elongation complexes with a sustained elongation block.

## Discussion

Genotoxic-induced transcriptional attenuation occurs in response to UV and gamma irradiation, as well as oxidative damage ^6–8,10^. However, the transcriptional response varies significantly depending on the nature of the damage. In contrast to previously described mechanisms of transcriptional attenuation following UV-induced damage, we find that oxidative damage induces a rapid and distinct transcriptional response, characterised by abrupt arrest of RNAPII within the gene body. Even more unexpectedly, we find that these RNAPII complexes restart transcription from their site of arrest. This is in contrast with the UV response, where transcriptional recovery relies on new transcription initiation and proceeds in 5’ to 3’ wavefront from the TSS ^7,10,13,47,48^. The difference most likely stems from the type of damage induced. While UV induced bulky DNA lesions act as roadblocks for RNAPII, and lesion stalled RNAPII itself plays a central role in recruiting of repair factors such as CSB ^5,9,13^, this is not the case following oxidative damage. Unlike the UV response, RNAPII is neither ubiquitinated and subsequently degraded, nor are NER factors such as XPC or CSB required for transcriptional restart. Indeed, high doses of hydrogen peroxide for which transcription fails to recover within a few hours, have been associated with RNAPII ubiquitination ^8,57^. We therefore speculate that RNAPII ubiquitination following oxidative stress serves only as a secondary response, facilitating removal of persistently stalled RNAPII from the DNA template. At lower doses of oxidative damage, RNAPII restarts from within the gene body, enabling a less disruptive and faster transcriptional recovery. Compared to the UV response, the kinetics of RNAPII arrest and restart following oxidative damage are extremely rapid. Whereas the UV-induced transcriptional repression is strongest a few hours after the initial exposure and recovery occurs 6-24 hours post-UV ^13^, oxidative damage-induced repression is immediate and recovery of overall transcription levels are restored within 1-2 hours of recovery. We suspect that this rapid response results from arrested RNAPII complexes being able to restart from within the gene body, thereby bypassing the need to physically remove RNAPII complexes from the DNA template. This aligns with our observation that RNAPII complexes arrested due to oxidative damage are differently configured in terms of their associated factors. Similarly to the heat shock response, we observed an accumulation of NELF-paused RNAPII during the transcriptional arrest and early recovery phase ^36^. However, unlike the heat shock response, oxidative stress did not result in NELF condensate formation, nor was the presence of NELF required for the transcriptional repression ^36^. In addition to its global role in heat shock-mediated transcriptional downregulation, NELF is involved in local transcriptional silencing through its recruitment to sites of DSBs in a PARylation-dependent manner ^28^. These observations led us to question whether NELF might be recruited to RNAPII within the gene body in response to oxidative stress. However, NELF ChIP-seq data excluded this possibility. Instead we find that NELF-mediated pausing is not per se required for the transcriptional response although it is impacting the pause site of RNAPII, leading to a transition of RNAPII pausing from the 1^st^ pause site to the 2^nd^ pause site (+1 nucleosome) ^51^. These observations again emphasise how global transcriptional repression can be achived through different mechanisms depending on the type of cellular stress and the nature of DNA damage induced.

Bulky DNA lesions induced by UV are known to block RNAPII progression both *in vitro* and *in vivo*, while non-bulky DNA lesions such as those typically induced by oxidative damage have generally been considered not to induce RNAPII stalling ^17–19^. SSBs however have been reported to stall RNAPII elongation *in vitro* ^21^. This is most likely not due to the DNA break itself, but rather to the aberrant 3’ termini of the gap impeding RNAPII progression ^58^. Our data strongly suggest that RNAPII gene body arrest is dependent on DNA damage, which is rapidly resolved through BER and SSBR, involving the DNA glycosylase OGG1 and XRCC1. XRCC1 serves as a scaffold protein to facilitate recruitment of DNA repair enzymes to SSB either generated directly enabling SSBR or to SSB generated as BER intermediates ^59^. Consequently, lack of XRCC1 leads to a delay in SSBR and is associated with neuronal dysfunction and neurological diseases, highlighting its important role in the maintenance of genome stability ^59,60^. However, the molecular interplay between SSBR and transcription has until now not been well-studied. Here we describe how the lack of XRCC1 leads to strong defects in RNAPII gene body recovery consistent with persistent occurrence of SSB within genes. Accordingly, cells lacking XRCC1 completely fail to recover transcription even at 4 hours after hydrogen peroxide treatment, as a consequence of sustained transcriptional repression within genes. Similar to XRCC1, OGG1i treatment results in very dramatic and prolonged gene body RNAPII arrest which is sustained at the 1-hour recovery timepoint. In contrast to lack of XRCC1, OGG1i-treated cells do eventually recover nascent transcription levels at the 2–4-hour recovery timepoint. We suspect this difference is due to additional DNA glycosylases such as NEIL1, NEIL2 and MUTYH ^61,62^. Indeed, NEIL1 and to a lesser extent NEIL2 have been shown to function as backup BER glycosylases working both on 8OG and further oxidation products such as hydantoin lesions upon OGG1 TH5487 inhibitor treatment or OGG1 depletion ^61^. This also fits well with mice knockout phenotypes, where individual BER glycosylase knockouts display only a moderate increase in mutation frequencies, while double or triple knockouts, such as OGG1/MUTYH knockouts display strong phenotypes characterised by high susceptibility to cancer and shortened lifespan ^63^. In addition, alternative repair pathways that might compensate for the lack of OGG1-mediated repair, such as NER, could facilitate a secondary response in case of persistently stalled RNAPII complexes ^44^. In contrast, XRCC1 is required both for SSBR and downstream in the BER pathway to resolve SSB intermediates ^59^. We speculate that this explains why lack of XRCC1 leads to sustained transcription repression, as lack of XRCC1 cannot easily compensated for by alternative repair pathways.

Oxidative damage-induced transcriptional repression coincides with high levels of PARylation which quickly decrease as transcription recovers. Sustained PARylation achieved through PARG inhibition maintains transcriptional repression, highlighting the dynamic regulation of the transcriptional response through PARylation. Traditionally, PARylation has been considered as a recruitment signal for DNA repair factors and plays a crucial role in SSBR through recruitment of XRCC1 ^23^. Lack of XRCC1 has also been suggested to promote PARP ‘trapping’ on DNA leading to a transcription block through maintained PARP1 binding to sites of DNA lesions ^56^. However, while we find that cells lacking XRCC1 fail to recover gene body transcription following oxidative stress, PARPi treatment exerts a milder effect on gene body transcription, suggesting that the presence of the oxidative DNA lesions themselves rather than PARylation-induced effects are the primary cause of gene body RNAPII arrest. Instead, we find that PARylation controls the release of early RNAPII complexes from the TSS already at the 15 min recovery timepoint. Notably, if we had relied solely on quantification of overall levels of nascent transcription following PARPi treatment, we would have concluded that PARPi treatment completely reverses the transcription inhibition as early as 15 min recovery. However, nascent transcriptomics clearly show that the effect of PARPi is more complex and spatially restricted to early RNAPII elongation complexes. Intriguingly, the effect of PARP inhibition of early RNAPII elongation complexes is independent of BER and SSBR DNA repair factors such as OGG1 or XRCC1, as we show that PARPi treatment ‘overrides’ continued transcriptional repression and promotes release of early elongation complexes even in cells lacking XRCC1 or following treatment with OGG1i. Based on our results, we suggest a spatial role of PARP1 which has hitherto been mistakenly overlooked.

The question remains how PARylation restricts early RNAPII elongation complexes. Both NELF and the pTEFb component CycT1 have been reported as direct PARylation targets ^14,64^. PARylation of NELF has been suggested to promote elongation ^64^, whereas CycT1 PARylation induced by MNNG (N-methyl-N’-nitro-N-nitrosoguanidine) disrupts CycT1 phase separation, correlating with reduced RNAPII hyperphosphorylation *in vitro* ^14^. In the case of CycT1, this would suggest an inhibitory effect on RNAPII pause release. However, it remains unknown if MNNG induces a spatially separated transcriptional response like hydrogen peroxide. Another possibility is that PARylation of proteins associated with early RNAPII complexes signals recruitment of additional factors such as NuMA which has been suggested to impact oxidative damage repair through a yet undefined mechanism ^55^. Thus, elucidation of the precise molecular mechanism of PARylation dependent restriction of early RNAPII elongation remains an exciting topic for future research and will undoubtedly provide new insight into how genotoxic DNA such as that induced by oxidative stress elicits a highly regulated transcriptional response.

As PARP inhibition is not able to prevent the initial repression of transcription, we suggest that the immediate and strong transcriptional attenuation reflects arrest of RNAPII by oxidative DNA lesions and RNAPII-blocking repair intermediates, occurring in a PARP-independent manner. This fits well with our results showing that the immediate transcriptional repression dramatically impacts RNAPII gene body activity. This is analogous to the situation for UV-induced bulky lesions, where the DNA damage itself directly impacts the transcription machinery. However, due to the type of DNA lesion and consequently how repair and RNAPII stalling are dealt with, the two types of genotoxic insults led to completely distinct outcomes in terms of transcriptional response. These differences are manifested in both the kinetics of the transcriptional response and factor dependencies, underscoring the unique nature of transcriptional regulation in response to oxidative damage.

While the coupling between transcription and nucleotide excision repair (NER) is well established, known as TC-NER, and relies on DNA damage recognition via RNAPII stalling within actively transcribed genes, the interplay between BER and transcription remains largely unexplored. A defining feature of TC-NER is its preferential repair of bulky DNA lesions on the transcribed strand of active genes ^7,9,13^. Our findings suggest that transcriptional recovery following oxidative damage is closely linked to DNA repair mediated by BER and SSBR, pointing to the possible existence of transcription-coupled BER (TC-BER) and transcription-coupled SSBR (TC-SSBR). It has been hypothesised that TC-BER may operate analogously to TC-NER, favoring repair within transcription units. While some studies support preferential BER of oxidative lesions in the transcribed strand in reconstituted transcription assays ^65^, others report no strand bias in the repair of 8-oxoguanine (8OG) lesions ^66^, leaving the mechanistic basis of such coupling unresolved. Unlike TC-NER, we do not observe direct recruitment of DNA repair factors to RNAPII. This absence may reflect a more transient interaction between the transcription machinery and repair complexes, or the inherently rapid kinetics of BER and SSBR compared to TC-NER ^13,59^. This underscores the need for further investigation into the nature and dynamics of transcription-coupled repair beyond NER.

In summary, our findings demonstrate that oxidative damage triggers a dramatic and immediate global repression of nascent transcription. This transcriptional attenuation is characterised by increased stalling of early RNAPII complexes near the TSS, as well as RNAPII arrest within the gene body. A distinctive feature of this response is that gene body–arrested RNAPII can resume transcription from within the gene body itself, enabling rapid recovery. Although NELF-mediated regulation influences promoter-proximal RNAPII pausing and positioning, NELF is not essential for the transcriptional repression. Strikingly, we find that separate mechanisms govern the transcriptional response of early RNAPII elongation complexes and gene body–arrested RNAPII. Early RNAPII stalling is regulated by PARylation, whereas recovery within the gene body depends on DNA repair mediated by BER and SSBR, suggesting that oxidative DNA lesions *in vivo* impede RNAPII progression. Thus, our temporally and spatially resolved characterisation of the nascent transcriptional response to oxidative damage reveals previously unrecognized layers of genotoxic-induced transcriptional regulation. These mechanisms serve to transiently repress global transcription through RNAPII arrest, orchestrated by PARylation and DNA damage repair pathways.

## Author contributions

L.H.G. and Q.A.T conceived the project. Q.A.T., L.H.G., L.W., E. L., H. K.M. I., S.K., D.L.M., and N.N.M. performed experiments. Q.A.T performed main experiments. L.H.G performed proof-of-concept experiments, assay establishment and initial TT_chem_-seq experiments. L.W. assisted in generating NELFC degron cell line, performed NELF microscopy experiments as well as UV and heat shock experiments. E.L. performed NELF and RNAPII ChIP-seq experiments. H.K.M.I. performed RNAPII Dsk2 pull-outs. S.K. performed initial EU assays. D.L.M performed *in vitro* RNAPII transcription assays. N.N.M. generated *XPC* knockout cell line. Q.A.T performed bioinformatic analysis, with input from H.L. L.H.G. wrote the manuscript with input from Q.A.T. All authors read and approved the final version of the manuscript.

## Acknowledgments

This work was supported by grants to L.H.G. from the European Research Council (ERC Agreement, TranscriptStress, 101076758), a Hallas-Møller Emerging Investigator Grant from the Novo Nordisk Foundation (NNF20OC0059959), and a Sapere Aude Research Leader Programme grant from the Independent Research Fund Denmark (0165-00092B). Research at the Center for Gene Expression (CGEN) is funded by the Danish National Research Foundation (DNRF166). Q.A.T. was supported by the Lundbeck Postdoctoral Fellowship Programme (LF R380-2021-1284). N.N.M. was supported by the EMBO Postdoctoral Fellowship (ALTF 911-2020). Mass spectrometry-based proteomics analyses were performed by the Proteomics Research Infrastructure (PRI) at the University of Copenhagen (UCPH), supported by the Novo Nordisk Foundation (grant agreement number NNF19SA0059305). We thank members of the CGEN/CPR/reNEW Sequencing Facility, University of Copenhagen, for expert technical assistance, and Jesper Q. Svejstrup for feedback on the manuscript.

## Methods

### Resource availability

Plasmids used in this study have been deposited in Addgene. Any additional information required to reanalyse the data reported in this paper is available from the lead contact upon request.

### Experimental Model

Flp-In^TM^ T-REx HEK293 cells (HEK293, R78007, ThermoFisher Scientific, human embryonic kidney epithelial, female origin) were cultured at 37°C with 5% CO_2_ in high glucose DMEM (Biowest, L0104) supplemented with 10% v/v FBS (Gibco, 10270106), 100 U/mL penicillin, 100 μg/mL streptomycin (Gibco, 10378016) 2 mM L-glutamine. Cells were routinely passaged 2-3 times a week. HEK293 CSB KO cells have previously been described ^67^. All cell lines were confirmed to be mycoplasma-free. Oxidative stress induction treatments were carried out for 15 min (or 30 min for EU-labelling) with direct addition of freshly prepare 1000x concentrate to cell culture media, fresh H_2_O_2_ (maximum 3 month old kept at 4°C) to final concentration of 1 mM H_2_O_2_ (Sigma, H1009) or freshly resuspended Menadione sodium bisulfite (MND, Sigma, M5750) in water to final concentration of 0.25 mM. After treatment the media was discarded, and cells were washed once with PBS pH = 7.4 (Gibco, 10010056) and replenished with fresh 37°C cell culture media. Inhibitors treatments were carried out by direct addition to the cell culture media with the following concentration 100 μM DRB (Sigma, D1916) or/and 10 μM Olaparib (PARPi, MedchemExpress, Cat#HY-10162) or/and 10 μM TH5487 (OGG1i, MedchemExpress, Cat#HY-125276) and 10 μM PDD 00017273 (PARGi, MedchemExpress, Cat#HY-108360). DRB inhibition was carried out 3.5 hours prior to other treatments. Inhibition of PARPi, PARGi and OGG1i were preformed 1 hour prior to other treatments unless specified otherwise. Cells treated with inhibitors were subjected to oxidative stress by direct addition of oxidative reagents to cell culture media. PARPi, OGG1i and PARGi inhibitors were then replenished together with cell culture media during stress recovery. DRB was either replenished or not depending on experiments. UV irradiation was performed with an exact dose of 20 J/m^2^ as in ^10^. Heat shock (HS) was performed by incubating cells at 43°C with pre-warmed media for 30 min or 1 hour.

### Plasmid construction and generation of stable cell lines Generation of stable cell lines

HEK293 Flp-In™ T-REx™ RPB3-3xFLAG cells (RPB3-FLAG) and U2OS Flp-In™ T-REx™ GFP-NELFE were generated by stable integration of the pDEST/TO/FRT/RPB3-3xFLAG or pFRT/TO/GFP-NELFE plasmids respectively. The human RPB3 (POLR2C) cDNA was amplified from pIRES-puro-RPB3-HA plasmid (gifted from Jesper Svejstrup) and NELFE cDNA were PCR amplified with Q5® High-Fidelity DNA Polymerase (NEB) from HEK293 WT cells genomic DNA extracted with Quick-DNA™ Miniprep Plus Kit (Zymo Research). Respective cDNA fragments were inserted through digestion and ligation into pENTR4 containing attB flanking sequences. RPB3 cDNA (in pENTR4-RPB3) and NELFE cDNA (in pENTR4-NELFE) were tagged through insertion into respectively pDEST/TO/3xFLAG and pDEST/GFP/NELFE destination vectors using Gateway™ LR Clonase II (Invitrogen, ThermoFisher Scientific), resulting with in-frame C-terminal fusion of RPB3 to 3xFLAG (3xDYKDDDDK) tag and N-terminal fusion of NELFE with EGFP. HEK293 Flp-In cells in DMEM without antibiotic complementation were transfected for 48 hours with 900 ng insertion plasmid pOG44 (ThermoFisher Scientific) and 100 ng donor plasmid pDEST/TO/RPB3-3xFLAG or pDEST/GFP/NELFE using Lipofectamine™ 3000 (Invitrogen, ThermoFisher Scientific; Cat#L3000015), following the manufacturer’s protocol. After 24 hours, the transfection cell culture media was replaced with fresh culture media for 24 hours, followed by one passage prior to selection with hygromycin (100 μg/mL), zeocin (100 μg/mL), and blasticidin (15 μg/mL) for 10 days. Single colonies were manually picked and passaged 2–3 times prior to testing. Tagged cell lines were treated with doxycycline (100 ng/mL) for 24 hours to induce RPB3-3xFLAG or GFP-NELFE confirmed by western blotting with respectively anti-FLAG (Sigma, Cat#F1804) and anti-GFP (Abcam, Cat#ab290) antibodies.

### Generation of knock-out cell lines

HEK293 Flp-In™ T-REx™ XRCC1 knockout cells (XRCC1 KO) were generated by CRISPR/Cas9-mediated genome editing. The pDG458-XRCC1/sg1&2 plasmid was constructed by cloning two guide RNAs (gRNAs: 5′-TGCAGGACACGACATGGCGG-3′ and 5′-AGCCCACGACGTTGACATGC-3′) into BbsI-digested pDG458 containing Cas9 fused to EGFP. This plasmid was transfected into HEK293 cells using Lipofectamine™ 3000 for 48 hours according to manufacturer’s protocol. GFP-positive cells were then sorted using a FACS system (FACSMelody™, BD Biosciences) into two 96-well plates. After 2–3 passages, individual clones were screened by western blotting to confirm XRCC1 knockout. Two validated knockout clones (KO#3 and KO#8) were used in subsequent experiments. HEK293 Flp-In™ T-REx™ XPC knockout cells (XPC KO) line was generated by CRISPR/Cas9-mediated genome editing. The gRNA sequence 5’-CAACATGGCTCGGAAACGCG-3’ was ligated into the vector pSpCas9(BB)-2A-Puro (pX459) V2.0 (Addgene, #48138) to generate the pX459_XPC_gRNA1 plasmid. HEK293 were transfected with the plasmid pX459_XPC_gRNA1 using Lipofectamine^TM^ 2000 for 48 hours following manufacturer instructions. Transfected cells were passaged and treated with puromycin (2 μg/mL) for 72 hours. Cells were then washed twice with PBS, detached with trypsin and quantified to prepare single cell dilutions in a 96 well plate. Two weeks after, colonies originated from a single cell were evaluated for XPC knock out by western blot using the anti-XPC antibody (D1M5Y, Cell Signalling, Cat#14768). Flp-In™ T-REx™ CSB knockout cells (CSB KO) were previously described.^10^

### Generation of NELFC degron cell line

HEK293 Flp-In™ T-REx FKBP12_F36V_-NELFCD cells (NELFCD-degron) were generated via CRISPR double break following homology recombination knock-in of a C-terminal FKBP12 degron tag, along with a cleavable puromycin resistance gene, into the endogenous HEK293 NELFCD locus with the donor plasmid pDONR-HA1/NELFCD/FKBP/PURO/HA2. To generate the donor plasmid, NELFCD cDNA sequence and homology arms (HA1 and HA2; >500 bp each) were amplified by PCR with Q5® High-Fidelity DNA Polymerase from HEK293 genomic DNA. The degron and puromycin resistance cassette were amplified from the pCRIS-PITChv2-Puro-dTAG (BRD4) plasmid (Addgene #91793). All fragments were assembled into a donor plasmid (pBluescript II SK(+)) using Gibson assembly with the NEBuilder® HiFi DNA Assembly Kit (NEB). To created double strand breaks and allow homology recombination targeted CRISPR components were delivered to cells, the pDG458-NELFCD-sg1&2 plasmid was constructed by cloning two gRNAs (5′-ATTTGCAGTGAGCTTTAACG-3′ and 5′-TGAAAGGGTTTTTCCACAAC-3′) into BbsI-digested pDG458 (Addgene, #100900). HEK293 cells were then co-transfected with pDONR-HA1/NELFCD/FKBP/PURO/HA2 and pDG458-NELFCD-sg1&2 using Lipofectamine^TM^ 3000 with manufacturer protocols in antibiotic-free cell culture media. After 24 hours, cells were selected with puromycin (2 μg/mL) for 48 hours. For selection of positive clones, colonies were picked after selection passaged 2-3 times and genomic DNA extracted with QuickExtract™ DNA Extraction Solution (Lucigen, Cat# LGCQE09050). A PCR fragment overlapping tagged region was amplified clones were selected based on fragment size shift and further confirmed by western blotting with NELFCD-specific antibodies. To further validate the degron system, homozygous knock-in clones were treated with 250 nM dTAG-v1 (Tocris Bioscience, Cat#6914) for 1 hour to induce *in vivo* NELFCD degradation. Depletion of NELFCD protein and reduction of chromatin-bound NELFA protein levels were confirmed by western blotting.

### EU-labelling assay

Approximately 7,000 HEK293 cells were seeded in 96-well Greiner CELLSTAR^®^ 96 well plate (ThermoFischer Scientific, Cat#M0562) coated for 15 min at room temperature with poly-D-Lysine (Gibco, Cat#A3890401) and washed twice with PBS. For treated samples, cell culture media was replaced by media complemented with indicated concentration of H_2_O_2_ or MND for 30 min and washed with PBS and replenished with new 37°C cell culture media. Nascent RNA was pulse-labelled for 30 min with addition of cell culture media complemented with 5-Ethynyluridine (EU, Jena Bioscience, Cat#CLK-N002, final concentration 750 μM). For treated samples (no recovery) both oxidative reagents and EU were added simultaneously. Labelled cells were immediately fixed with 3.7 % formaldehyde diluted in PBS for 45 min in the dark. The cells were then washed three times in PBS and permeabilized with 0.5 % Triton X-100 in PBS for 30 min at room temperature in the dark. Nascent RNA was then visualized by click-it chemistry, the EU-incorporated RNA was labelled in the dark for 2 hours with a mix of 5 μM Alexa Fluor^TM^ 488 Azid (ThermoFischer Scientific, Cat#A10266), 4 mM copper sulfate (Sigma-Aldrich) and 100 mM ascorbic acid (Sigma-Aldrich). Cells were then washed twice with PBS and incubated for 30 min at room temperature in the dark with 2 μg/mL DAPI (Sigma-Aldrich, Cat#D9742) diluted in PBS. Cells were washed twice with PBS and kept in the dark at 4°C in PBS for up to 48 hours. A total of 25 images per well were acquired with the ScanR acquisition system (Olympus) controlling a motorized Olympus IX-81 wide-field microscope and analysed and quantified with the ScanR Analysis software.

### Total RNA extraction

Total RNA was extracted by adding TRIzol™ Reagent (ThermoFisher Scientific, Cat#15596026) directly to the cell monolayer after cell culture media removal. Chloroform was added to the cell/TRIzol mixture at a 1:5 volume ratio, followed by thorough mixing for 30 seconds and centrifugation at 12,000 g for 15 min at 4 °C. The upper aqueous phase was collected and further purified by adding 1 volume of chloroform/isoamyl alcohol (24:1; Sigma-Aldrich, Cat#C0549), followed by vortexing and centrifugation at 12,000 × g for 5 min at 4 °C. The resulting upper phase, containing total RNA, was precipitated by adding one volume of isopropanol (Sigma-Aldrich, Cat#I9516) and 30 ng/mL GlycoBlue™ (Invitrogen, ThermoFisher Scientific, Cat#AM9515) and incubated at room temperature for 20 min. The RNA was pelleted by centrifugation at 20,000 g for 30 min and washed once with ice-cold 85% ethanol. The final RNA pellet was resuspended in RNase-free water.

### Nascent RNA dot blot

Dot blots were used to detect global levels of nascent transcription based on 4-thiorudine (4sU, Glentham Life Sciences, Cat#GN6085) RNA incorporation. Dot blot were performed as described in ^38^ with minor changes. Approximately 500,000 cells were seeded in 6-well plates. The next day, 4sU (final concentration 1mM) was added directly to the tissue culture for 15 min coordinated with H_2_O_2_ treatment and recovery time points, labelling was then arrested with addition of 0.5 mL TRIzol. Total RNA was extracted and 3 μg of 4sU labelled total RNA was biotinylated for 2 hours in the dark with 50 μL of 0.1 mg/ml EZ-link^TM^ HPDP-Biotin (ThermoFischer Scientific, Cat#21341 in DMF) with 3 μL of biotin buffer (833 mM Tris-HCl pH 7.4, and 83.3 mM EDTA) for a final reaction volume of 300 μL. Unreacted biotin-linker was removed from RNA samples by addition of 1.1 volumes of ROTI^®^Aqua-P/C/I (Roth, Cat#985.1) vortexed, precipitated with 1.1 isopropanol followed by pellet centrifugation, ethanol washes and resuspension in RNAse-free H_2_O. Purified RNA sample were resuspended in 10 μL and applied to a Hybond-N+ membrane (Cytiva, Cat#RPN203B) in a dot blot apparatus (BioRad). The immobilized RNA was UV-crosslinked to the membrane with UV-C 0.24 J/cm^2^ and the membrane incubated in blocking solution (PBS pH = 7.4; 10% SDS; 1 mM EDTA) for 20 min at room temperature followed by at 15 min incubation with a 1:50,000 dilution of 1 mg/mL streptavidin-horseradish peroxidase (HRP) (ThermoFisher Scientific Cat#N100) in blocking solution. The membrane was then washed twice in blocking solution buffer for 10 min, followed by two washes in dot blot wash buffer I (PBS pH = 7.4; 1% SDS) for 10 min each and two washes in dot blot wash buffer II (PBS pH = 7.4; 0.1% SDS) for 10 min each. The biotin-bound HRP was visualized with 2 min incubation of 1:4 diluted ECL detection reagent (BioRad) on an ChemiDoc Imaging System (BioRad). As a loading control, the membrane was stained with methylene blue solution (0.5 M sodium acetate; 0.5 % methylene blue) for 10 min, then de-stained by several washes in water.

### TT_chem_-seq

TT_chem_-seq was performed as described previously with minor modifications ^38^. In brief, 10 cm dishes with 80% confluent cells RNA were labelled *in vivo* for 15 min with 4sU by adding it directly to the tissue culture media to a final concentration of 1 mM. Labelling was terminated by addition of TRIzol and total RNA was extracted by TRIzol/chloroform extraction followed by isopropanol precipitation. Per samples 80-100 μg of total RNA resuspended in 100 μL RNAse-free H_2_O was complemented with 1 % (relative to total RNA concentration) of *S. cerevisiae* 4tU-labelled RNA spike-in. Total RNA was fragmented by adding 20 μL of freshly prepared 1 M NaOH per sample, thoroughly mixed and incubated for 40 min on ice to < 400 nt fragments. RNA fragmentation was terminated by addition of 80 μL 1 M Tris-HCl pH = 6.8 and samples were immediately purified with 1 volume of ROTI^®^Aqua-P/C/I (ROTH, Cat# X985.1) extraction and precipitated with isopropanol. Resuspended and purified 4sU labelled RNA in RNAse-free water was biotinylated in 10 mM Tris-HCl pH = 7.4; 1 mM EDTA complemented with 0.5 mg/ μL MTSEA biotin-XX linker (Biotum, Cat# BT90066) for 30 min in the dark at room temperature. Biotinylated RNA was immediately purified by ROTI^®^Aqua-P/C/I extraction and isopropanol precipitation and resuspended in 50 μL RNAse-free water. To enrich for 4sU labelled RNA, the purified RNA was denatured at 65°C for 10 min, with 5 min on ice was used, and isolated with μMACS Streptavidin Kit (Miltenyi, Cat#130-074-101). A total of 100uL μMACS Streptavidin MicroBeads were added to denatured 4sU total RNA and incubated for 15 min at room temperature with agitation. Beads bound to 4sU biotinylated RNA were applied to magnetic μColumns and transferred to the magnetic field of the μMACS magnetic separator (Miltenyi). The columns were washed twice with 1 mL 55°C pre-warmed pull-down wash buffer (100 mM Tris-HCl pH 7.4, 10 mM EDTA, 1 M NaCl and 0.1 % (v/v) Tween 20) and eluted with two consecutive washes with freshly prepared 100 mM DTT. Biotinylated RNA was further cleaned up from DTT elution by ROTI^®^Aqua-P/C/I extraction and isopropanol precipitation and resuspended in RNAse-free H_2_O. Concentration of 4sU enriched RNA was measured by Qubit (ThermoFischer Scientific, Cat# Q32852) using the RNA HS assay kit and size of fragmented RNA assessed by TapeStation using RNA ScreenTape Assay for TapeStation Systems (Agilent, Cat# 5067-5576). 30 ng of purified 4sU biotinylated RNA was used to prepare libraries with NEBNext® Ultra™ II RNA library prep kit (NEB, Cat#E7760) according to manufacturer’s protocol (protocol 5 for fragmented RNA) with ligation of two barcodes of 8 nt and 11 nt UMIs (NEB, Cat#E7416). Libraries were amplified using 8 cycles of PCR and sequenced as single-end 68 bp reads on a NextSeq 2000 platform (Illumina).

### Cell growth assays

Approximately 5,000 HEK293 cells per well were seeded in triplicates in 96-well plates, pre-coated with poly-D-Lysine (Gibco) for 15 min and washed twice with PBS. The following day cell treatments were performed in 96-wells by addition of cell culture media complemented with different concentration of H_2_O_2_ or MND for 15 min. The complemented media was removed, and cells were washed once with PBS and fresh 37°C cell culture media was replenishment. Growth was monitored using an IncuCyte S3 Live-Cell Analysis System (Sartorius). Images of live cells were captured every 2 hours for up to 6 days. Images were analysed in the IncuCuyte S3 image analysis software, with an output of cell confluency (%).

### Cell fractionation and RNAPII immunoprecipitation

Cell pellets from one 15 cm plate were resuspended in 1 mL volume HE buffer (10 mM HEPES-NaOH pH=7.5; 10 mM KCl, protease and phosphatase inhibitors: Vanadate, Sodium fluoride, and ß-glycerolphosphate) and incubated 15 min on ice. Nuclei were pelleted 15 min; 1,000 g and the supernatant taken as the cytoplamic fraction. The nuclear pellet was resuspended in 500 ml NE buffer (20 mM HEPES-NaOH pH = 7.9; 150 mM NaCl; 0.05% NP40; 10% glycerol; protease and phosphatase inhibitors fresh). Chromatin were pelleted 15 min; 20,000 g and the supernatant taken as the nucleoplasmic fraction. Cytoplasmic and nucleoplasmic fractions were pooled as soluble fraction. Chromatin pellets were digested in 500 mL of CD Buffer (20 mM HEPES-NaOH pH=7.9; 150 mM NaCl; 1.5 mM MgCl_2_; 0.05% NP40; 10% glycerol; protease and phosphatase inhibitors fresh; 250 U/ml DENARASE^®^ (c-Lecta, Cat#20804) and incubated on a rotating wheel for 1 hour at 4°C to digest nucleic acids and solubilize chromatin and release chromatin-bound proteins. Digested chromatins were pelleted 15 min at 20,000 g and the supernatant was taken as the enriched chromatin fraction. For immunoprecipitation, Dynabeads^TM^ Protein G Magnetic Beads (Invitrogen, ThermoFischer Scientific, Cat#10004D) were washed and resuspended in 2 beads volumes of PBS + 0.02% Tween 20. The resuspended beads were conjugated for 1 hour at room temperature with 1 mg 4H8 or 3E10 as indicated (for capture of phosphorylated RNAPII complexes) or with anti-FLAG (Sigma, #1806, for capture of total RNAPII complexes in a RPB3-FLAG expressing cell line). Non-antibody conjugated beads were used for pull-out as negative control for mass spectrometry. Following antibody incubation, unconjugated antibodies were removed by from beads by two washes with PBS + 0.02 % (v/v) Tween20. Protein concentrations of chromatin enriched fractions were measured with Protein Assay Dye Reagent (BioRad, Cat#5000001) through absorbance measured at 495 nm and concentration was adjusted to the sample with the lowest concentration by addition of CD buffer (constituting the ‘input’ sample for western blotting). Concentration-adjusted chromatin fractions were added to antibody-conjugated beads (or unconjugated beads for negative controls) and incubated for 3 hours at 4°C with agitation. Samples were placed on a magnetic rack and the supernatant was removed and kept (‘unbound’ lysate for western blotting). Proteins bound to magnetic beads were then washed three times with IP wash buffer (10 mM Tris-HCl pH=7.5; 150 mM NaCl; 0.05% NP40; 0.5 mM EDTA; protease and phosphatase inhibitors fresh). Beads bound proteins were either extracted by boiling with 2xSDS sample buffer (10 mM Tris-HCl; pH = 6.8, 4% SDS, 0.2% bromophenol blue, 20% Glycerol, 200 mM DTT) for western blotting (‘IP’ sample) or for mass spectrometry analysis, the beads were washed two additional times with TBS and stored dry at -20°C until further processing for proteomics.

### Mass spectrometry

All samples for proteomic analysis were carried out in triplicate and samples were prepared in parallel from the point of cell seeding. TBS washed frozen beads were thawed and incubated for 30 min with elution buffer 1 (50 mM Tris-HCl pH = 7.5; 2 M Urea; 2 mM DTT; 20 μg/ml trypsin) followed by a second elution for 5 min with elution buffer 2 (50 mM Tris-HCl pH = 7.5; 2 M Urea; 10 mM Chloroacetamide). Both eluates were combined and further incubated at room temperature over-night. Tryptic peptide mixtures were acidified to 1% TFA and loaded on Evotips (Evosep). Peptides were separated on 15 cm, 150 μM ID columns packed with C18 beads (1.9 μM) (Pepsep) on an Evosep ONE HPLC applying the ‘30 samples per day’ method and injected via a CaptiveSpray source and 10 μm emitter into a timsTOF pro mass spectrometer (Bruker) operated in PASEF mode.

### Western blotting

For whole cell extracts, cell pellets were collected, resuspended in whole cell extraction buffer (20 mM Tris-HCl pH = 7.5; 300 mM NaCl; 2.5 mM MgCl2; 1% NP40; 10% glycerol) with freshly added DENARASE^®^ (c-Lecta, Cat#20804), fresh protease and phosphatase inhibitors, and then incubated rotating at 4°C for 1 hour followed by centrifugation at 10,000 g for 2 min at 4°C. Supernatant was transferred to a new tube and protein concentration measured using Bio-Rad protein assay reagent. For whole cell extracts, chromatin enriched fractions and IP fractions, protein concentrations were adjusted to the sample with the lowest concentration with the respective appropriate extraction buffer. 10 μL protein samples was complemented with 10 μL 2x SDS sample buffer and boiled at 98°C for 5 min. 20 μL protein sample were separated by SDS-PAGE gel electrophoresis (6%, 8%, 10% or 15% acrylamide were used) in running buffer (25 mM Tris pH = 8.3, 192 mM glycine, 0.1% SDS,) and transferred to nitrocellulose membranes by wet transfer in transfer buffer (0.025 M Tris, 0.192 M glycine, 10% (v/v) ethanol). Membranes were stained with Ponceau S solution for 10 min and washed with dH_2_O, and a picture was acquired. Membranes were then blocked in blocking buffer (5% (w/v) milk in TBS-T) and incubated with primary antibody (in 5% (w/v) milk in TBS-T) overnight at 4°C.

Membranes were then washed with TBS-T, and incubated with HRP-conjugated secondary antibodies in blocking buffer for 1 hour at room temperature and then washed several times in TBS-T. HRP signal detection was obtained using 1:1 ratio of each Clarity ECL Western Blotting Substrates (BioRad, Cat#1705061) applied for 2min onto the membrane and visualization by ChemiDoc Imaging System (BioRad).

### ELCAP-seq

ELCAP-seq was carried out similarly to previously described ^39^. RPB3-FLAG cells were seeded in 4x15 cm dished per samples; RPB3-FLAG was induced by adding a final concentration of 100 ng/mL doxycycline for 16 hours directly to cell culture media. Induced cells were treated with 1 mM H_2_O_2_ for 15 min or let recover after a PBS wash and replenishment of warm cell culture media for variable recovery times. Immediately cells were collected, centrifuged for 2 min at 1,000 g and the pellets frozen in liquid nitrogen. Frozen cells were then immediately used for cell fractionation as described above. A volume of 25 μL Dynabeads^TM^ Protein G Magnetic Beads per sample were washed and conjugated to 1 mg anti-FLAG (Sigma, #1806) as previously described in immunoprecipitation section. Protein concentrations from chromatin enriched fraction were measured with Protein Assay Dye Reagent (BioRad) and adjusted to the sample with the lowest concentration with CD buffer. Normalised chromatin enriched fractions were then added to FLAG-conjugated magnetic beads for 3 hours at 4°C with agitation. The beads were washed 4 times with ELCAP wash buffer (10 mM Tris-HCl pH = 7.5; 250 mM NaCl; 0.5% (v/v) NP-40; 0.5 mM EDTA; fresh protease inhibitor; protease Inhibitors cocktail and fresh phosphatase inhibitors). Short RNA fragments protected by RNAPII were extracted with the Quick-RNA MicroPrep kit (Zymo, Cat#R1050) according to the manufacture’s instruction for short RNA extraction (17-200 nt). Briefly, RNPAII bound beads were incubated for 5 min at RT in extraction buffer (per 100 μL sample; 100 μL of supplied Zymo RNA lysis Buffer, 100 μL of IP wash buffer,100 μL ethanol). RNA was isolated using kit supplied Zymo-Spin™ IC columns following the manufacturer’s instructions. The size distribution of extracted RNA was assessed with the Bioanalyzer Small RNA Analysis system (Agilent). Library were prepared using 30 ng of short RNAs with NEBNext® Small RNA library preparation kit (NEB, Cat# E7300) according to the manufacturer’s protocol with the only modification being inclusion of a UMI containing adapter; the initial ligation was carried with the custom ligation adapter ‘5’-App-NNNNNNAGATCGGAAGAGCACACGTCT-3’ (IDT) with the same ligation conditions as manufacturer’s protocol. Libraries were amplified with 8 cycles of PCR and the whole library was separated by size with Novex™ TBE Gels, 6% (Invitrogen, ThermoFischer Scientific, Cat# EC6265BOX), stained with SYBR^TM^gold (Invitrogen, ThermoFischer Scientific, Cat# S11494) and fragment size ranging from 147 nt to 190 nt were excised from the gel and purified according to NEBNext® Small RNA library preparation kit protocol. Size selected libraries were then sequenced as single-end run (68 bp reads) on a NextSeq2000 platform (Illumina).

### Ubiquitinated proteins pull-down

DSK2 beads were prepared as previously described ^40^ with minor changes. In brief *E. coli* expressing DSK2 were diluted in ice cold dilution buffer (1x PBS, 0.5% Triton X-100, protease inhibitors) and sonicated (15 s ON, 30 s OFF, total 10 min ON time). HEK293 cells were either treated with UV (20 J/m^2^) and let recover for 45 min after cell culture media replenishment or with 1 mM H_2_O_2_ for 15 min and recovery time as indicated. Cell pellets were resuspended in TENT buffer (50 mM Tris-HCl pH = 7.4; 2 mM EDTA; 150 NaCl; 1 % Triton X-100; 2,5 µM MgCl; protease and phosphatase inhibitors added fresh and 250 U/ml DENARASE®) and incubated 1 hour at 4°C. Debris and non-solubilized proteins were removed by centrifugation at 14,000 g for 10 min. Protein concentration of the resulting supernatant was determined with Protein Assay Dye Reagent (Biorad). Lysates protein concentration was equalized before adding 25 µL DSK2 beads and incubated overnight at 4°C. DSK2 beads enriched for ubiquitin bound proteins were spun down at 700 g for 5 min and washed once in TENT buffer and once with PBS (pH = 7.4). To release ubiquitinated proteins from the beads, samples were heated at 95°C for 5 min in 2x SDS sample buffer. The samples were separated from the beads by centrifugation and 1% input and 20 % sample were used for western blotting.

### Chromatin immunoprecipitation and sequencing (ChIP-seq)

For each sample, 80 % confluent HEK293 cells in a 15 cm plate were crosslinked with 1% formaldehyde for 15 min and stopped with 125 mM glycine for 5 min. Fixed cells were then homogenized in cellular lysis buffer (5 mM Pipes pH = 8; 85 mM KCl; 0.5 % NP-40). Nuclei were then resuspended in nuclear lysis buffer (50 mM Tris pH = 8.1; 10 mM EDTA; 1 % SDS) and sonicated to obtain DNA fragments between 50 bp and 500 bp (5 cycles; 30sec ON, 30sec OFF, with Bioruptor pico). Samples were diluted 10 times in dilution buffer (0.01 % SDS; 1.1 % Triton X-100; 1.2 mM EDTA; 16.7 mM Tris pH = 8; 167 mM NaCl). Samples were pre-cleared with 25 µL proteins A/G beads (ThermoFisher Scientific, Cat#20422) and blocked with 500 µg BSA for each 200 µg of chromatin for 2 hours at 4°C. A volume of 100 µL pre-cleared chromatin was keep at -20°C for inputs and another volume of chromatin was then incubated overnight with either NELFE or total RPB1 (D8L4Y) antibodies at 4°C. Immunoprecipitation was performed using 35 µL proteins A/G beads (blocked with 500 µg BSA) for 2 hours at 4°C. Beads were then washed once with dialysis buffer (2 mM EDTA; 50 mM Tris-HCl pH=8.1; 0.2% Sarkosyl), once wash buffer (100 mM Tris pH = 8.8; 500 mM LiCl; 1% NP-40; 1% NaDoc) and once with TE (10 mM Tris pH = 8; 0.5 mM EDTA). Immunoprecipitated complexes and inputs were resuspended in TE and incubated with 1 µL of RNase A (10 mg/mL, ThermoFisher Scientific, Cat#EN0531) for 30 min at 37°C. Reverse crosslinking was performed by incubating samples overnight at 70°C with 4 µL SDS (10%) and incubation with ProteinaseK (10 mg/mL, ThermoFisher Scientifc Cat#AM2548) for 1 hour 30 min at 45°C. DNA was purified by phenol/chlorophorm extraction followed by ethanol precipitation. Libraries were prepared with NEBNext® Ultra™ II DNA library prep kit (NEB, Cat#E7604) with 12 nt UMIs adaptors (NEB, Cat#E7395) according to manufacturer’s protocol and sequenced on a NextSeq2000 platform (Illumina).

### *In vitro* transcription with pig thymus RNAPII

RNAPII was purified from pig thymus as previously described ^68^. The elution of the 8WG16 conjugated beads was dialyzed in dialyze buffer (20 mM HEPES-NaOH pH = 8; 150 mM NaCl; 10 % Glycerol; 2 mM β-mercaptoethanol) and concentrated with Amicon^®^ Ultra 4 centrifugal filter (50 kDa) (Millipore, Cat#UF8010). The elongation complex was assembled as previously described with minor modification ^69^. Briefly, 5 pmol of pre-annealed RNA:DNA (template strand) hybrid was mixed with an equimolar amount of pure RNAPII and incubated 20 min at 30°C, followed by the addition of 10 pmol 5’ biotin-labelled non-template strand DNA for an additional 10 min. The assembled elongation complex was diluted with transcription buffer (20 mM Tris-HCl pH= 7.5; 100 mM NaCl; 8 mM MgCl2; 10 μM ZnCl2; 10% glycerol). The transcription reaction was started by addition of 1 mM rNTPs for 2 min at 25°C and stopped with 20 mM EDTA, 0.5 μL Proteinase K and 45% formamide. For treatment, H_2_O_2_ was added before the rNTPs at the indicated concentrations. Samples were incubated for 30 min at 50°C followed by 10 min at 70°C. RNA products were resolved in 8 M Urea, 16% polyacrylamide, 1x TBE gel. Gels were scanned in a Typhoon scanner (Cytiva) to detect Cy2 fluorescence.

### Immunofluorescence microscopy

GFP-tagged NELFE U2OS cells were seeded (with 100 ng/mL doxycycline) at 60-70% confluency onto coverslips coated with Poly-D-Lysine (Gibco) in 12-well plates. The next day cells were treated for 15 min or 30 min with 1 mM H_2_O_2_ and for 30 min or 1 hour heat shock at 43°C. Cells were then fixed with 4% (v/v) formaldehyde for 15 min at room temperature, washed once with PBS, and mounted onto slides using VECTASHIELD® Antifade Mounting Medium with DAPI (VWR, Cat#VECTH-1200). GFP and DAPI fluorescence were visualised with an inverted confocal microscope (Zeiss LSM 900)

### Image analysis

For EU-labelling experiments, scanR software was used to quantify each isolated nuclei green signal relative to DAPI signal. For picture plotting, raw tiff images were upload to ImageJ’s Fiji, untreated samples (HEK293 or *XRCC1* KO cells) were corrected for brightness and contrast; the same correction was then applied to all other images of the same cell line.

### TT_chem_-seq analysis

Raw fastq files for Read 1 were labelled with appropriate UMI reads using UMItools ^70^(v1.1.4) extract function followed by adapter trimming with cutadapt (v4.5) within trim_galore (v0.6.10) and read length filtering > 19nt. For the purposes of visualization, biological replicates UMI and adapter trimmed reads fastq files were merged and processed further similarly to individual replicates. In the case of *XRCC1* KO, two individual clones (KO#3 and KO#8) were considered as biological duplicates. Merged and individual samples were mapped with STAR^71^ (v2.7.11b) against a merged human-yeast genome assemblies (*Homo sapiens* GRCh38 and *Saccharomyces cerevisiae* sacCer, Ensembl release 108) with options as default. Genome alignments BAM files were filtered for multimapped reads with samtools (v1.21) view function (-q 10). UMI duplicates were then removed with UMItools dedup function. A yeast gene-level count matrix was generated with Subread (v2.0.6) featureCounts^72^ and used to calculate yeast size-factors with DEseq2^73^ (v1.42.0) “estimateSizeFactors” function. Strand specific and spike-in normalised BigWig files were generated from strand-specific bam files using Deeptools^74^ (v3.5.5) bamCoverage function (--scaleFactor = 1/ yeast size-factor) with bin size of 25 nt (-bs 25). TT_chem_-seq scaled plots (metagene profiles or heatmaps) were created from spike-in normalised BigWig files with DeepTools computeMatrix function (scale-regions, between –2 kb TSS to TES + 2 kb, bin size 50bp) using gene definitions from Ensembl (release 108). All gene features were selected for protein coding genes non-overlapping (extended to 5 kb upstream the TSS and 5 kb downstream of the TES) other coding genes (n = 11,593). ‘Long Genes’ derived from the later and were filtered with length > 90 kb (n = 2,217). Unscaled (reference-point) mode metagene profiles were centred on TSS with various distance around the TSS. TT_chem_-seq signal relative to untreated control ratio BigWig files were created using the DeepTools bigwigCompare function; the genome was partitioned to 100 bp bins for scaled metagene profiles and 50 bp bins for unscaled metagene profiles. Heatmaps were sorted by gene length (decreasing or increasing). Quantitative differential expression analysis of TT_chem_-seq data for Supplementary Fig. 2d was done through DESeq2 with replacement of human estimateSizeFactor by yeast spike-in size-factors. For Supplementary Fig. 5b BigWig files were converted to bedGraph, forward and reverse strand were then merged into one bedGraph file. In R (v4.3.2), with a custom script, coverage values were extracted from each feature with regards to strand and normalised to region size. Genomic spike-in normalised track screenshots were exported from IGB as vector images.

### ELCAP-seq analysis

Raw fastq files were trimmed with cutadapt (v4.5) within trim_galore (v0.6.10) and minimum read length filtering > 19nt and maximum read length < 47 nt. For purposes of visualization, biological replicates adapter trimmed and size filtered reads fastq files were merged and processed further similarly to individual replicates. Resulting reads were mapped with STAR with (–-alignEndsType Extend5pOfRead1) against the human genome assembly (GRCh38, Ensembl release 108). Genome alignment BAM files were filtered for multimapped reads and separated by strand with samtools view function (-q 10, -F 16 or -f 16). Single nucleotide strand specific and RPKM (read per million kilobase) normalised BigWig files were generated using Deeptools’ bamCoverage function for each stranded BAM files in single nucleotide resolution mode with the following options “-bs 1 --Offset -1 --normalizeUsing RPKM”. ‘Pause sites’ were obtained by calculating the highest peak position within pausing region (-20 nt , TSS, +120 nt) in the untreated sample with CAGEfightR^75^ .

ELCAP-seq centred metagene plots (profiles or heatmaps) were created using deepTools computeMatrix in reference-point mode (centred on TSS or the ‘pause sites’) over the same non-overlapping protein coding gene set as for TT_chem_-seq plots. ELCAP-seq signal relative to untreated control ratio BigWig files were created using the deepTools’ bigwigCompare function in single nucleotide mode (-bs 1). Heatmaps were sorted by gene length (decreasing or increasing). For Supplementary Fig. 5c ratio between TT_chem_-seq and ELCAP-seq BigWig files for each time points were created using deepTools bigwigCompare function and scaled to a normalising ratio calculated for library size dispersion between the two datasets. For Supplematery Fig. 5d ELCAP-seq BigWig files were converted to bedGraph, forward and reverse strand were then merged into one bedGraph file. In R, with a custom script, the pausing index (PI) was calculated as ratio of coverage values from TSS (TSS + 200 bp) over coverage for gene body region (TSS + 200 bp to TES – 200 bp) relative to size. The recurrence of pausing index was then plotted as distribution.

### Mass spectrometry analysis

Raw mass spectrometry data were analysed with MaxQuant (v1.6.15.0). Peak lists were searched against the human Uniprot FASTA database combined with 262 common contaminants by the integrated Andromeda search engine. False discovery rate was set to 1 % for both peptides (minimum length of 7 amino acids) and proteins. “Match between runs” (MBR) was enabled with a Match time window of 0.7, and a Match ion mobility window of 0.05 min. Relative protein amounts were determined by the MaxLFQ algorithm with a minimum ratio count of two. Statistical analysis of LFQ derived protein expression data was performed using the automated analysis pipeline of the Clinical Knowledge Graph.93 Protein entries referring to potential contaminants, proteins identified by matches to the decoy reverse database, and proteins identified only by modified sites, were removed. LFQ intensity values were normalised by log2 transformation and proteins with less than 70 % of valid values in at least one group were filtered out. The remaining missing values were imputed using the MinProb approach (random draws from a Gaussian distribution; width = 0.2 and downshift = 1.8) or with Mixed imputation using KNN.

### ChIP-seq analysis

NELFE, total RPB1 and input raw fastq files were trimmed with with trim_galore (paired-end mode) with minimum read length filtering > 19 nt. Filtered and trimmed paired end reads were mapped with STAR against human assembly GRCh38 (Ensembl release 108). Genome alignment BAM files were filtered for multi-mapped reads with samtools view function (-q 10). Bam files for NELFE or total RPB1 samples were normalised to input with MACS^76^ (v2.2.9.1). Resulting bedGraph files were used to compute heatmaps. In Fig. 4e input normalised NELFE ChIP-seq signal levels were used to sort (in decreasing order) both bottom panels of ELCAP-seq signals. In Fig. 4f ChIP-seq ratio (sample relative to untreated control) BigWig files were created using the deepTools bigwigCompare function, the genome was partitioned to 50 bp bins. NELFE ChIP-seq ratio BigWig files were then normalised to total RPB1 ChiP-seq levels with bigwigCompare function and plotted relative to NELFE ChIP-seq decreasing signal.

## References

1. Tubbs, A., and Nussenzweig, A. (2017). Endogenous DNA Damage as a Source of Genomic Instability in Cancer. Cell 168, 644–656. 10.1016/j.cell.2017.01.002.

2. Crow, J.F. (2000). The origins, patterns and implications of human spontaneous mutation. Nat Rev Genet 1, 40–47. 10.1038/35049558.

3. Lodato, M.A., Rodin, R.E., Bohrson, C.L., Coulter, M.E., Barton, A.R., Kwon, M., Sherman, M.A., Vitzthum, C.M., Luquette, L.J., Yandava, C.N., et al. (2018). Aging and neurodegeneration are associated with increased mutations in single human neurons. Science 359, 555–559. 10.1126/science.aao4426.

4. Caron, P., van der Linden, J., and van Attikum, H. (2019). Bon voyage: A transcriptional journey around DNA breaks. DNA Repair (Amst) 82, 102686. 10.1016/j.dnarep.2019.102686.

5. Lans, H., Hoeijmakers, J.H.J., Vermeulen, W., and Marteijn, J.A. (2019). The DNA damage response to transcription stress. Nat Rev Mol Cell Biol 20, 766–784. 10.1038/s41580-019-0169-4.

6. Wang, J., Muste Sadurni, M., and Saponaro, M. (2023). RNAPII response to transcription-blocking DNA lesions in mammalian cells. FEBS J 290, 4382–4394. 10.1111/febs.16561.

7. Gregersen, L.H., and Svejstrup, J.Q. (2018). The Cellular Response to Transcription-Blocking DNA Damage. Trends Biochem Sci 43, 327–341. 10.1016/j.tibs.2018.02.010.

8. Heine, G.F., Horwitz, A.A., and Parvin, J.D. (2008). Multiple mechanisms contribute to inhibit transcription in response to DNA damage. J Biol Chem 283, 9555–9561. 10.1074/jbc.M707700200.

9. van der Meer, P.J., and Luijsterburg, M.S. (2025). The molecular basis of human transcription-coupled DNA repair. Nat Cell Biol 27, 1230–1239. 10.1038/s41556-025-01715-9.

10. Tufegdzic Vidakovic, A., Mitter, R., Kelly, G.P., Neumann, M., Harreman, M., Rodriguez-Martinez, M., Herlihy, A., Weems, J.C., Boeing, S., Encheva, V., et al. (2020). Regulation of the RNAPII Pool Is Integral to the DNA Damage Response. Cell 180, 1245–1261 e1221. 10.1016/j.cell.2020.02.009.

11. Nakazawa, Y., Hara, Y., Oka, Y., Komine, O., van den Heuvel, D., Guo, C., Daigaku, Y., Isono, M., He, Y., Shimada, M., et al. (2020). Ubiquitination of DNA Damage-Stalled RNAPII Promotes Transcription-Coupled Repair. Cell 180, 1228–1244 e1224. 10.1016/j.cell.2020.02.010.

12. Mayne, L.V., and Lehmann, A.R. (1982). Failure of RNA synthesis to recover after UV irradiation: an early defect in cells from individuals with Cockayne’s syndrome and xeroderma pigmentosum. Cancer Res 42, 1473–1478.

13. Nieto Moreno, N., Olthof, A.M., and Svejstrup, J.Q. (2023). Transcription-Coupled Nucleotide Excision Repair and the Transcriptional Response to UV-Induced DNA Damage. Annu Rev Biochem 92, 81–113. 10.1146/annurev-biochem-052621-091205.

14. Fu, H., Liu, R., Jia, Z., Li, R., Zhu, F., Zhu, W., Shao, Y., Jin, Y., Xue, Y., Huang, J., et al. (2022). Poly(ADP-ribosylation) of P-TEFb by PARP1 disrupts phase separation to inhibit global transcription after DNA damage. Nat Cell Biol 24, 513–525. 10.1038/s41556-022-00872-5.

15. Adamowicz, M., Hailstone, R., Demin, A.A., Komulainen, E., Hanzlikova, H., Brazina, J., Gautam, A., Wells, S.E., and Caldecott, K.W. (2021). XRCC1 protects transcription from toxic PARP1 activity during DNA base excision repair. Nat Cell Biol 23, 1287–1298. 10.1038/s41556-021-00792-w.

16. Lindahl, T. (1993). Instability and decay of the primary structure of DNA. Nature 362, 709–715. 10.1038/362709a0.

17. Charlet-Berguerand, N., Feuerhahn, S., Kong, S.E., Ziserman, H., Conaway, J.W., Conaway, R., and Egly, J.M. (2006). RNA polymerase II bypass of oxidative DNA damage is regulated by transcription elongation factors. EMBO J 25, 5481–5491. 10.1038/sj.emboj.7601403.

18. Damsma, G.E., and Cramer, P. (2009). Molecular basis of transcriptional mutagenesis at 8-oxoguanine. J Biol Chem 284, 31658–31663. 10.1074/jbc.M109.022764.

19. Tornaletti, S., Maeda, L.S., Kolodner, R.D., and Hanawalt, P.C. (2004). Effect of 8-oxoguanine on transcription elongation by T7 RNA polymerase and mammalian RNA polymerase II. DNA Repair (Amst) 3, 483–494. 10.1016/j.dnarep.2004.01.003.

20. Oh, J., Fleming, A.M., Xu, J., Chong, J., Burrows, C.J., and Wang, D. (2020). RNA polymerase II stalls on oxidative DNA damage via a torsion-latch mechanism involving lone pair-pi and CH-pi interactions. Proc Natl Acad Sci U S A 117, 9338–9348. 10.1073/pnas.1919904117.

21. Kathe, S.D., Shen, G.P., and Wallace, S.S. (2004). Single-stranded breaks in DNA but not oxidative DNA base damages block transcriptional elongation by RNA polymerase II in HeLa cell nuclear extracts. J Biol Chem 279, 18511–18520. 10.1074/jbc.M313598200.

22. Jackson, S.P., and Bartek, J. (2009). The DNA-damage response in human biology and disease. Nature 461, 1071–1078. 10.1038/nature08467.

23. Ray Chaudhuri, A., and Nussenzweig, A. (2017). The multifaceted roles of PARP1 in DNA repair and chromatin remodelling. Nat Rev Mol Cell Biol 18, 610–621. 10.1038/nrm.2017.53.

24. Shanbhag, N.M., Rafalska-Metcalf, I.U., Balane-Bolivar, C., Janicki, S.M., and Greenberg, R.A. (2010). ATM-dependent chromatin changes silence transcription in cis to DNA double-strand breaks. Cell 141, 970–981. 10.1016/j.cell.2010.04.038.

25. Kakarougkas, A., Ismail, A., Chambers, A.L., Riballo, E., Herbert, A.D., Kunzel, J., Lobrich, M., Jeggo, P.A., and Downs, J.A. (2014). Requirement for PBAF in transcriptional repression and repair at DNA breaks in actively transcribed regions of chromatin. Mol Cell 55, 723–732. 10.1016/j.molcel.2014.06.028.

26. Dong, C., West, K.L., Tan, X.Y., Li, J., Ishibashi, T., Yu, C.H., Sy, S.M.H., Leung, J.W.C., and Huen, M.S.Y. (2020). Screen identifies DYRK1B network as mediator of transcription repression on damaged chromatin. Proc Natl Acad Sci U S A 117, 17019–17030. 10.1073/pnas.2002193117.

27. Chou, D.M., Adamson, B., Dephoure, N.E., Tan, X., Nottke, A.C., Hurov, K.E., Gygi, S.P., Colaiacovo, M.P., and Elledge, S.J. (2010). A chromatin localization screen reveals poly (ADP ribose)-regulated recruitment of the repressive polycomb and NuRD complexes to sites of DNA damage. Proc Natl Acad Sci U S A 107, 18475–18480. 10.1073/pnas.1012946107.

28. Awwad, S.W., Abu-Zhayia, E.R., Guttmann-Raviv, N., and Ayoub, N. (2017). NELF-E is recruited to DNA double-strand break sites to promote transcriptional repression and repair. EMBO Rep 18, 745–764. 10.15252/embr.201643191.

29. El-Khamisy, S.F., Masutani, M., Suzuki, H., and Caldecott, K.W. (2003). A requirement for PARP-1 for the assembly or stability of XRCC1 nuclear foci at sites of oxidative DNA damage. Nucleic Acids Res 31, 5526–5533. 10.1093/nar/gkg761.

30. Hanzlikova, H., Gittens, W., Krejcikova, K., Zeng, Z., and Caldecott, K.W. (2017). Overlapping roles for PARP1 and PARP2 in the recruitment of endogenous XRCC1 and PNKP into oxidized chromatin. Nucleic Acids Res 45, 2546–2557. 10.1093/nar/gkw1246.

31. Aprile-Garcia, F., Tomar, P., Hummel, B., Khavaran, A., and Sawarkar, R. (2019). Nascent-protein ubiquitination is required for heat shock-induced gene downregulation in human cells. Nat Struct Mol Biol 26, 137–146. 10.1038/s41594-018-0182-x.

32. Cavadas, M.A.S., Cheong, A., and Taylor, C.T. (2017). The regulation of transcriptional repression in hypoxia. Exp Cell Res 356, 173–181. 10.1016/j.yexcr.2017.02.024.

33. Mahat, D.B., Salamanca, H.H., Duarte, F.M., Danko, C.G., and Lis, J.T. (2016). Mammalian Heat Shock Response and Mechanisms Underlying Its Genome-wide Transcriptional Regulation. Mol Cell 62, 63–78. 10.1016/j.molcel.2016.02.025.

34. Vihervaara, A., Duarte, F.M., and Lis, J.T. (2018). Molecular mechanisms driving transcriptional stress responses. Nat Rev Genet 19, 385–397. 10.1038/s41576-018-0001-6.

35. Gressel, S., Schwalb, B., and Cramer, P. (2019). The pause-initiation limit restricts transcription activation in human cells. Nat Commun 10, 3603. 10.1038/s41467-019-11536-8.

36. Rawat, P., Boehning, M., Hummel, B., Aprile-Garcia, F., Pandit, A.S., Eisenhardt, N., Khavaran, A., Niskanen, E., Vos, S.M., Palvimo, J.J., et al. (2021). Stress-induced nuclear condensation of NELF drives transcriptional downregulation. Mol Cell. 10.1016/j.molcel.2021.01.016.

37. Cugusi, S., Mitter, R., Kelly, G.P., Walker, J., Han, Z., Pisano, P., Wierer, M., Stewart, A., and Svejstrup, J.Q. (2022). Heat shock induces premature transcript termination and reconfigured the human transcriptome. Mol Cell 82, 1573–1588 e1510. 10.1016/j.molcel.2022.01.007.

38. Gregersen, L.H., Mitter, R., and Svejstrup, J.Q. (2020). Using TTchem-seq for profiling nascent transcription and measuring transcript elongation. Nat Protoc 15, 604–627. 10.1038/s41596-019-0262-3.

39. Gregersen, L.H., Mitter, R., and Svejstrup, J.Q. (2022). Elongation factor-specific capture of RNA polymerase II complexes. Cell Rep Methods 2, 100368. 10.1016/j.crmeth.2022.100368.

40. Tufegdzic Vidakovic, A., Harreman, M., Dirac-Svejstrup, A.B., Boeing, S., Roy, A., Encheva, V., Neumann, M., Wilson, M., Snijders, A.P., and Svejstrup, J.Q. (2019). Analysis of RNA polymerase II ubiquitylation and proteasomal degradation. Methods 159-160, 146–156. 10.1016/j.ymeth.2019.02.005.

41. Gonzalo-Hansen, C., Steurer, B., Janssens, R.C., Zhou, D., van Sluis, M., Lans, H., and Marteijn, J.A. (2024). Differential processing of RNA polymerase II at DNA damage correlates with transcription-coupled repair syndrome severity. Nucleic Acids Res 52, 9596–9612. 10.1093/nar/gkae618.

42. Guo, J., Hanawalt, P.C., and Spivak, G. (2013). Comet-FISH with strand-specific probes reveals transcription-coupled repair of 8-oxoGuanine in human cells. Nucleic Acids Res 41, 7700–7712. 10.1093/nar/gkt524.

43. Kyng, K.J., May, A., Brosh, R.M., Jr., Cheng, W.H., Chen, C., Becker, K.G., and Bohr, V.A. (2003). The transcriptional response after oxidative stress is defective in Cockayne syndrome group B cells. Oncogene 22, 1135–1149. 10.1038/sj.onc.1206187.

44. Ranes, M., Boeing, S., Wang, Y., Wienholz, F., Menoni, H., Walker, J., Encheva, V., Chakravarty, P., Mari, P.O., Stewart, A., et al. (2016). A ubiquitylation site in Cockayne syndrome B required for repair of oxidative DNA damage, but not for transcription-coupled nucleotide excision repair. Nucleic Acids Res 44, 5246–5255. 10.1093/nar/gkw216.

45. Tuo, J., Jaruga, P., Rodriguez, H., Dizdaroglu, M., and Bohr, V.A. (2002). The cockayne syndrome group B gene product is involved in cellular repair of 8-hydroxyadenine in DNA. J Biol Chem 277, 30832–30837. 10.1074/jbc.M204814200.

46. Andrade-Lima, L.C., Veloso, A., Paulsen, M.T., Menck, C.F., and Ljungman, M. (2015). DNA repair and recovery of RNA synthesis following exposure to ultraviolet light are delayed in long genes. Nucleic Acids Res 43, 2744–2756. 10.1093/nar/gkv148.

47. Williamson, L., Saponaro, M., Boeing, S., East, P., Mitter, R., Kantidakis, T., Kelly, G.P., Lobley, A., Walker, J., Spencer-Dene, B., et al. (2017). UV Irradiation Induces a Non-coding RNA that Functionally Opposes the Protein Encoded by the Same Gene. Cell 168, 843–855 e813. 10.1016/j.cell.2017.01.019.

48. Lavigne, M.D., Konstantopoulos, D., Ntakou-Zamplara, K.Z., Liakos, A., and Fousteri, M. (2017). Global unleashing of transcription elongation waves in response to genotoxic stress restricts somatic mutation rate. Nat Commun 8, 2076. 10.1038/s41467-017-02145-4.

49. Zumer, K., Maier, K.C., Farnung, L., Jaeger, M.G., Rus, P., Winter, G., and Cramer, P. (2021). Two distinct mechanisms of RNA polymerase II elongation stimulation in vivo. Mol Cell 81, 3096–3109 e3098. 10.1016/j.molcel.2021.05.028.

50. Stein, C.B., Field, A.R., Mimoso, C.A., Zhao, C., Huang, K.L., Wagner, E.J., and Adelman, K. (2022). Integrator endonuclease drives promoter-proximal termination at all RNA polymerase II-transcribed loci. Mol Cell 82, 4232–4245 e4211. 10.1016/j.molcel.2022.10.004.

51. Aoi, Y., Smith, E.R., Shah, A.P., Rendleman, E.J., Marshall, S.A., Woodfin, A.R., Chen, F.X., Shiekhattar, R., and Shilatifard, A. (2020). NELF Regulates a Promoter-Proximal Step Distinct from RNA Pol II Pause-Release. Mol Cell 78, 261–274 e265. 10.1016/j.molcel.2020.02.014.

52. DeBerardine, M., Booth, G.T., Versluis, P.P., and Lis, J.T. (2023). The NELF pausing checkpoint mediates the functional divergence of Cdk9. Nat Commun 14, 2762. 10.1038/s41467-023-38359-y.

53. Bai, P. (2015). Biology of Poly(ADP-Ribose) Polymerases: The Factotums of Cell Maintenance. Mol Cell 58, 947–958. 10.1016/j.molcel.2015.01.034.

54. Slade, D., Dunstan, M.S., Barkauskaite, E., Weston, R., Lafite, P., Dixon, N., Ahel, M., Leys, D., and Ahel, I. (2011). The structure and catalytic mechanism of a poly(ADP-ribose) glycohydrolase. Nature 477, 616–620. 10.1038/nature10404.

55. Ray, S., Abugable, A.A., Parker, J., Liversidge, K., Palminha, N.M., Liao, C., Acosta-Martin, A.E., Souza, C.D.S., Jurga, M., Sudbery, I., and El-Khamisy, S.F. (2022). A mechanism for oxidative damage repair at gene regulatory elements. Nature 609, 1038–1047. 10.1038/s41586-022-05217-8.

56. Demin, A.A., Hirota, K., Tsuda, M., Adamowicz, M., Hailstone, R., Brazina, J., Gittens, W., Kalasova, I., Shao, Z., Zha, S., et al. (2021). XRCC1 prevents toxic PARP1 trapping during DNA base excision repair. Mol Cell 81, 3018–3030 e3015. 10.1016/j.molcel.2021.05.009.

57. Inukai, N., Yamaguchi, Y., Kuraoka, I., Yamada, T., Kamijo, S., Kato, J., Tanaka, K., and Handa, H. (2004). A novel hydrogen peroxide-induced phosphorylation and ubiquitination pathway leading to RNA polymerase II proteolysis. J Biol Chem 279, 8190–8195. 10.1074/jbc.M311412200.

58. Neil, A.J., Belotserkovskii, B.P., and Hanawalt, P.C. (2012). Transcription blockage by bulky end termini at single-strand breaks in the DNA template: differential effects of 5’ and 3’ adducts. Biochemistry 51, 8964–8970. 10.1021/bi301240y.

59. Caldecott, K.W. (2024). Causes and consequences of DNA single-strand breaks. Trends Biochem Sci 49, 68–78. 10.1016/j.tibs.2023.11.001.

60. Hoch, N.C., Hanzlikova, H., Rulten, S.L., Tetreault, M., Komulainen, E., Ju, L., Hornyak, P., Zeng, Z., Gittens, W., Rey, S.A., et al. (2017). XRCC1 mutation is associated with PARP1 hyperactivation and cerebellar ataxia. Nature 541, 87–91. 10.1038/nature20790.

61. Hanna, B.M.F., Michel, M., Helleday, T., and Mortusewicz, O. (2021). NEIL1 and NEIL2 Are Recruited as Potential Backup for OGG1 upon OGG1 Depletion or Inhibition by TH5487. Int J Mol Sci 22. 10.3390/ijms22094542.

62. David, S.S., O’Shea, V.L., and Kundu, S. (2007). Base-excision repair of oxidative DNA damage. Nature 447, 941–950. 10.1038/nature05978.

63. Xie, Y., Yang, H., Cunanan, C., Okamoto, K., Shibata, D., Pan, J., Barnes, D.E., Lindahl, T., McIlhatton, M., Fishel, R., and Miller, J.H. (2004). Deficiencies in mouse Myh and Ogg1 result in tumor predisposition and G to T mutations in codon 12 of the K-ras oncogene in lung tumors. Cancer Res 64, 3096–3102. 10.1158/0008-5472.can-03-3834.

64. Gibson, B.A., Zhang, Y., Jiang, H., Hussey, K.M., Shrimp, J.H., Lin, H., Schwede, F., Yu, Y., and Kraus, W.L. (2016). Chemical genetic discovery of PARP targets reveals a role for PARP-1 in transcription elongation. Science 353, 45–50. 10.1126/science.aaf7865.

65. Banerjee, D., Mandal, S.M., Das, A., Hegde, M.L., Das, S., Bhakat, K.K., Boldogh, I., Sarkar, P.S., Mitra, S., and Hazra, T.K. (2011). Preferential repair of oxidized base damage in the transcribed genes of mammalian cells. J Biol Chem 286, 6006–6016. 10.1074/jbc.M110.198796.

66. Thorslund, T., Sunesen, M., Bohr, V.A., and Stevnsner, T. (2002). Repair of 8-oxoG is slower in endogenous nuclear genes than in mitochondrial DNA and is without strand bias. DNA Repair (Amst) 1, 261–273. 10.1016/s1568-7864(02)00003-4.

67. Mulderrig, L., Garaycoechea, J.I., Tuong, Z.K., Millington, C.L., Dingler, F.A., Ferdinand, J.R., Gaul, L., Tadross, J.A., Arends, M.J., O’Rahilly, S., et al. (2021). Aldehyde-driven transcriptional stress triggers an anorexic DNA damage response. Nature 600, 158–163. 10.1038/s41586-021-04133-7.

68. Hu, X., Malik, S., Negroiu, C.C., Hubbard, K., Velalar, C.N., Hampton, B., Grosu, D., Catalano, J., Roeder, R.G., and Gnatt, A. (2006). A Mediator-responsive form of metazoan RNA polymerase II. Proc Natl Acad Sci U S A 103, 9506–9511. 10.1073/pnas.0603702103.

69. Han, Z., Moore, G.A., Mitter, R., Lopez Martinez, D., Wan, L., Dirac Svejstrup, A.B., Rueda, D.S., and Svejstrup, J.Q. (2023). DNA-directed termination of RNA polymerase II transcription. Molecular Cell. 10.1016/j.molcel.2023.08.007.

70. Smith, T., Heger, A., and Sudbery, I. (2017). UMI-tools: modeling sequencing errors in Unique Molecular Identifiers to improve quantification accuracy. Genome Res 27, 491–499. 10.1101/gr.209601.116.

71. Dobin, A., Davis, C.A., Schlesinger, F., Drenkow, J., Zaleski, C., Jha, S., Batut, P., Chaisson, M., and Gingeras, T.R. (2013). STAR: ultrafast universal RNA-seq aligner. Bioinformatics 29, 15–21. 10.1093/bioinformatics/bts635.

72. Liao, Y., Smyth, G.K., and Shi, W. (2014). featureCounts: an efficient general purpose program for assigning sequence reads to genomic features. Bioinformatics 30, 923–930. 10.1093/bioinformatics/btt656.

73. Love, M.I., Huber, W., and Anders, S. (2014). Moderated estimation of fold change and dispersion for RNA-seq data with DESeq2. Genome Biol 15, 550. 10.1186/s13059-014-0550-8.

74. Ramirez, F., Dundar, F., Diehl, S., Gruning, B.A., and Manke, T. (2014). deepTools: a flexible platform for exploring deep-sequencing data. Nucleic Acids Res 42, W187–191. 10.1093/nar/gku365.

75. Thodberg, M., Thieffry, A., Vitting-Seerup, K., Andersson, R., and Sandelin, A. (2019). CAGEfightR: analysis of 5’-end data using R/Bioconductor. BMC Bioinformatics 20, 487. 10.1186/s12859-019-3029-5.

76. Zhang, Y., Liu, T., Meyer, C.A., Eeckhoute, J., Johnson, D.S., Bernstein, B.E., Nusbaum, C., Myers, R.M., Brown, M., Li, W., and Liu, X.S. (2008). Model-based analysis of ChIP-Seq (MACS). Genome Biol 9, R137. 10.1186/gb-2008-9-9-r137.

