## Supplemental Figures for "Oxidative stress triggers RNAPII arrest through PARylation and DNA damage"

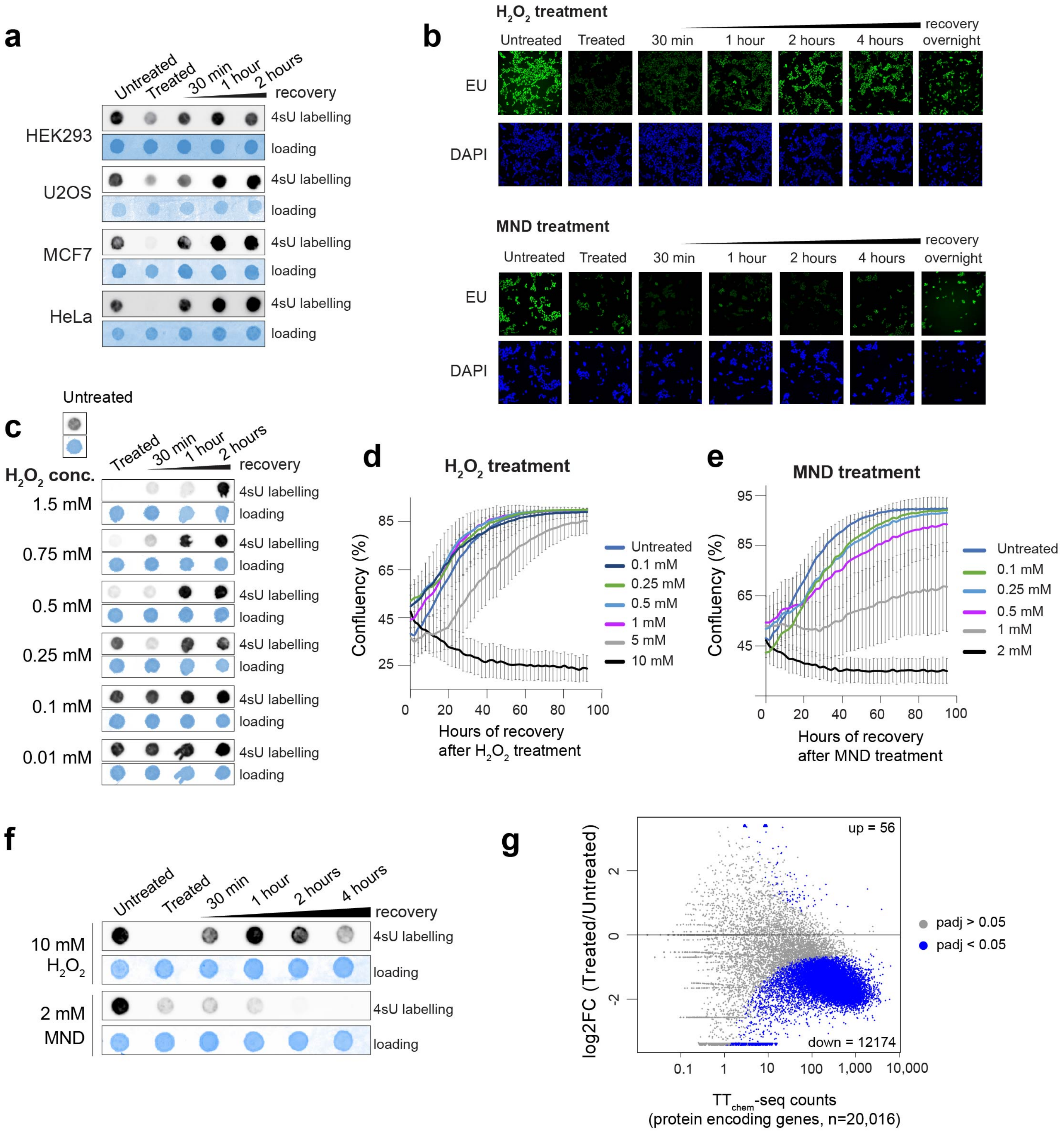


**Supplementary Fig. 1**

**Supplementary Fig. 1. Oxidative stress represses nascent transcription. a** Nascent transcription levels (4sU dot blot) in HEK293, U2OS, MCF7 and HeLa cells treated for 15 with 1 mM $H_2O_2$ and following media replenishment. Staining with methylene blue serves as a loading control. **b** Representative images of global nascent transcription (EU immunofluorescence) from experiments in Figure 1c and 1d of WT HEK293 cells following oxidative stress conditions. All image intensities are displayed with same exposure as the untreated control. **c** Nascent RNA levels (4sU dot blot) of cells treated with decreasing $H_2O_2$ doses for 15 min. Staining with methylene blue serves as a loading control. **d-e** Growth assay of HEK293 cells monitored by live cell imaging (IncuCyte). Cells were treated for 15 min with indicated doses of $H_2O_2$ (d) or MND (e), followed by media replenishment. **f** Nascent RNA levels (4sU dot blot) cells treated for 15 min with 10 mM $H_2O_2$ and 2 mM MND with indicated recovery times after media replenishment. Staining with methylene blue serves as a loading control. **g** Scatterplot of log2 fold-change between the mean $TT_{chem}$-seq normalised (with spike-in) read counts per protein encoding-genes of three biological replicates of $H_2O_2$ treated cells compared to untreated cells, relative to mean read counts for all protein coding genes (n = 20,016). Blue dots represent genes with a false discovery rate (FDR) corrected q-value < 0.05 measured with Wald test.

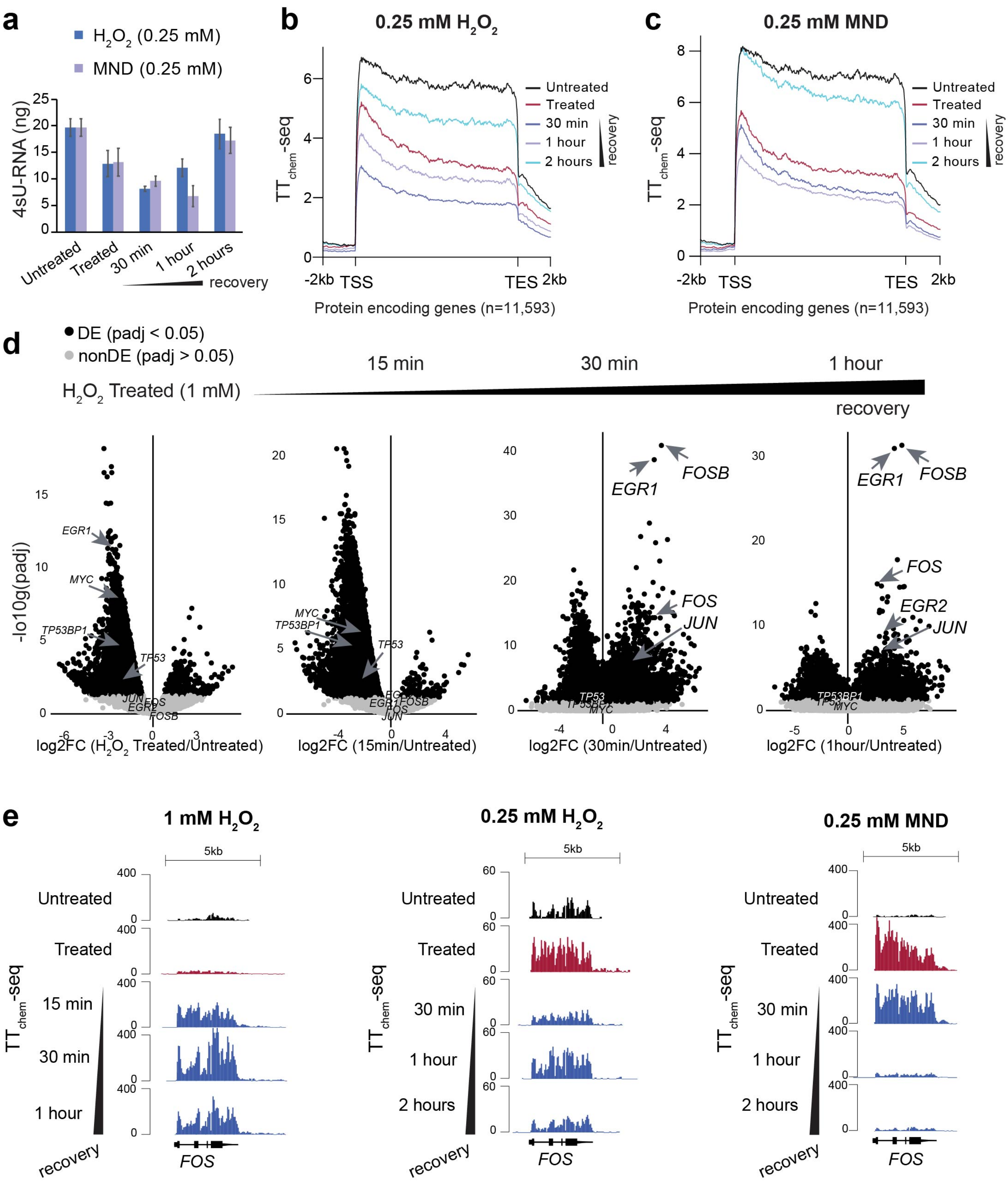


**Supplementary Fig. 2**

**Supplementary Fig. 2. The transcriptional response to varying doses of ROS. a** 4sU labelled RNA yield measured after streptavidin pull-down for duplicates of untreated HEK293 cells, treated for 15 min with 0.25 mM $H_2O_2$ or 0.25 mM MND and indicated recovery times. **b-c** Metagene analysis of $TT_{chem}$-seq data for non-overlapping coding genes (n=11,593) of untreated and indicated recovery timepoint following 0.25 mM $H_2O_2$ treatment of HEK293 cells. (b) or 0.25 mM MND. (c) Lines is the mean of spike-in normalised read counts between two biological replicates. **d** Scatterplots of fdr p-adjusted value (-log10 padj) of comparison between the means of triplicates $TT_{chem}$-seq spike-in normalised signals of $H_2O_2$ treated or recovery cells compared to untreated cells, relative to log2 fold-change for all annotated genomic features (n = 60,649). Genes in black have Wald test padj < 0.05 and in grey padj > 0.05. Immediate early genes *EGR1*, *EGR2*, *FOS*, *FOSB*, *TP53*, *TP53BP1* and *MYC* are highlighted. **e** Single-gene view of $TT_{chem}$-seq data for the immediate early stress response genes *FOS* for experimental conditions in Figure 3a and S2a-c.

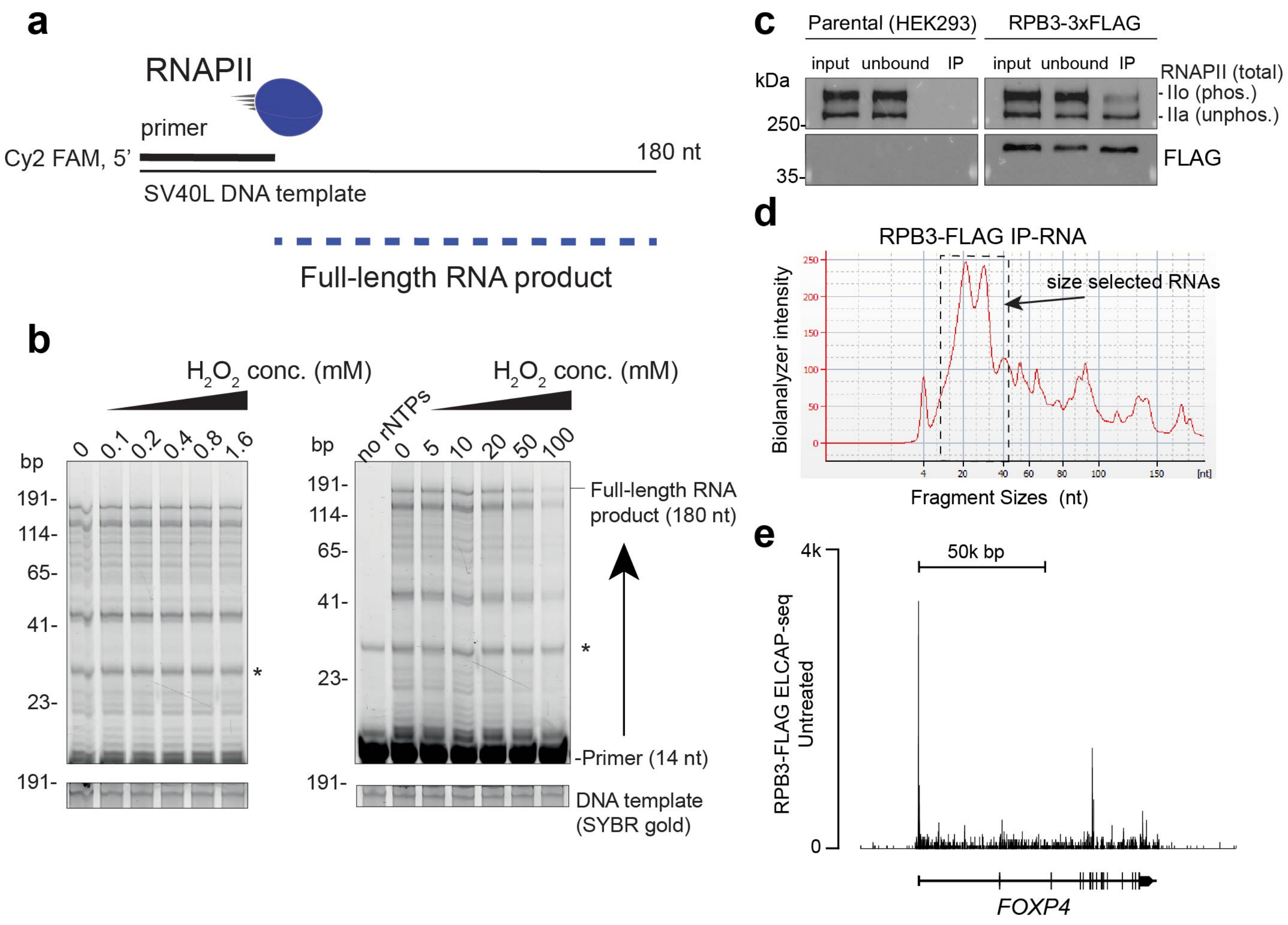


**Supplementary Fig. 3**

**Supplementary Fig. 3.** ***In vitro*** **RNAPII activity after ROS exposure and validation of FLAG-RBP3 expressing cell line for total RNAPII ELCAP-seq. a** Setup used for RNAPII *in vitro* transcription assay. RNAPII complexes purified from pig thymus is loaded on the FAM-labelled primer bound to SV40L DNA template resulting in various sized RNA fragments up to 180 nt. **b** Reconstituted *in vitro* transcription assay performed with RNAPII complexes treated with indicated $H_2O_2$ concentrations for 15 min. The resulting transcripts from the *in vitro* transcription assay were resolved on 8 M Urea TBE gels and visualised by Cy2 fluorescence. **c** Verification of RNAPII IP used in the ELCAP protocol from HEK293-RPB3-FLAG cell line. Western blot analysis of total RNAPII using either a RBP1 antibody (D8L4Y) or FLAG antibody. Input, unbound, and IP fractions were resolved. **d** Bioanalyzer spectrum of RNA extracted from RPB3-FLAG IPs and used as input for ELCAP-seq. Detected RNA intensity is shown for different size fragments (nt). Dotted line represents the region size-selected for library preparation. **e** Single-gene view of ELCAP-seq signal (TPM) for the *FOXP4* gene in untreated conditions.

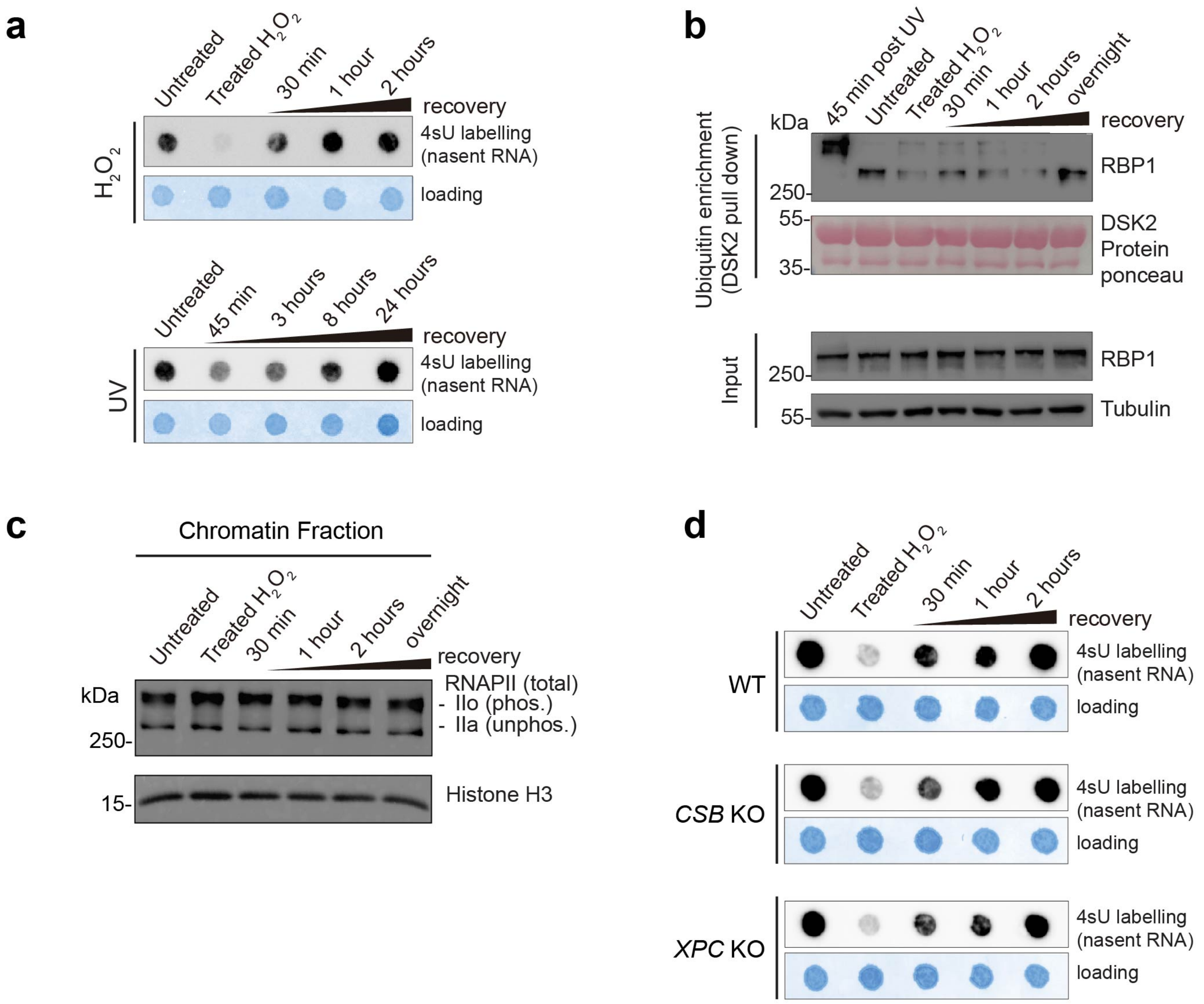


**Supplementary Fig 4**

**Supplementary Fig. 4. The transcriptional response to oxidative stress is distinct from the UV-induced transcriptional response and not dependent on NER factors CSB or XPC. a** Nascent transcription levels (4sU dot blot) in HEK293 cells treated with 1 mM $H_2O_2$ at indicated times after media washout or recovery time following 20 J/m$^2$ UV-C (256 nm) irradiation. Staining with methylene blue serves as a loading control. **b** Western blot of phosphorylated RNAPII (4H8) in DSK2-enriched proteins fractions (ubiquitin enriched) for HEK293 cells 45 min after 20 J/m$^2$ UV-C or 1 mM $H_2O_2$ treatment and indicated times after media replenishment. Ponceau represents DSK2 protein loading control (top). Input fraction was probed with a pan RPB1 phosphorylated antibody (4H8) and tubulin for loading control (bottom). **c-d** Western blot of total RNAPII (D8L4Y) level of chromatin-enriched cellar fraction after 1 mM $H_2O_2$ treatment and indicated recovery time after media replenishment. (Same experiment as shown in Figure 1I). Histone H3 serves as a loading control. **d** Nascent transcription levels (4sU dot blot) in WT (HEK293), *CSB* KO and *XPC* KO cells treated with 1 mM $H_2O_2$ and indicated times after media replenishment. Staining with methylene blue serves as a loading control.

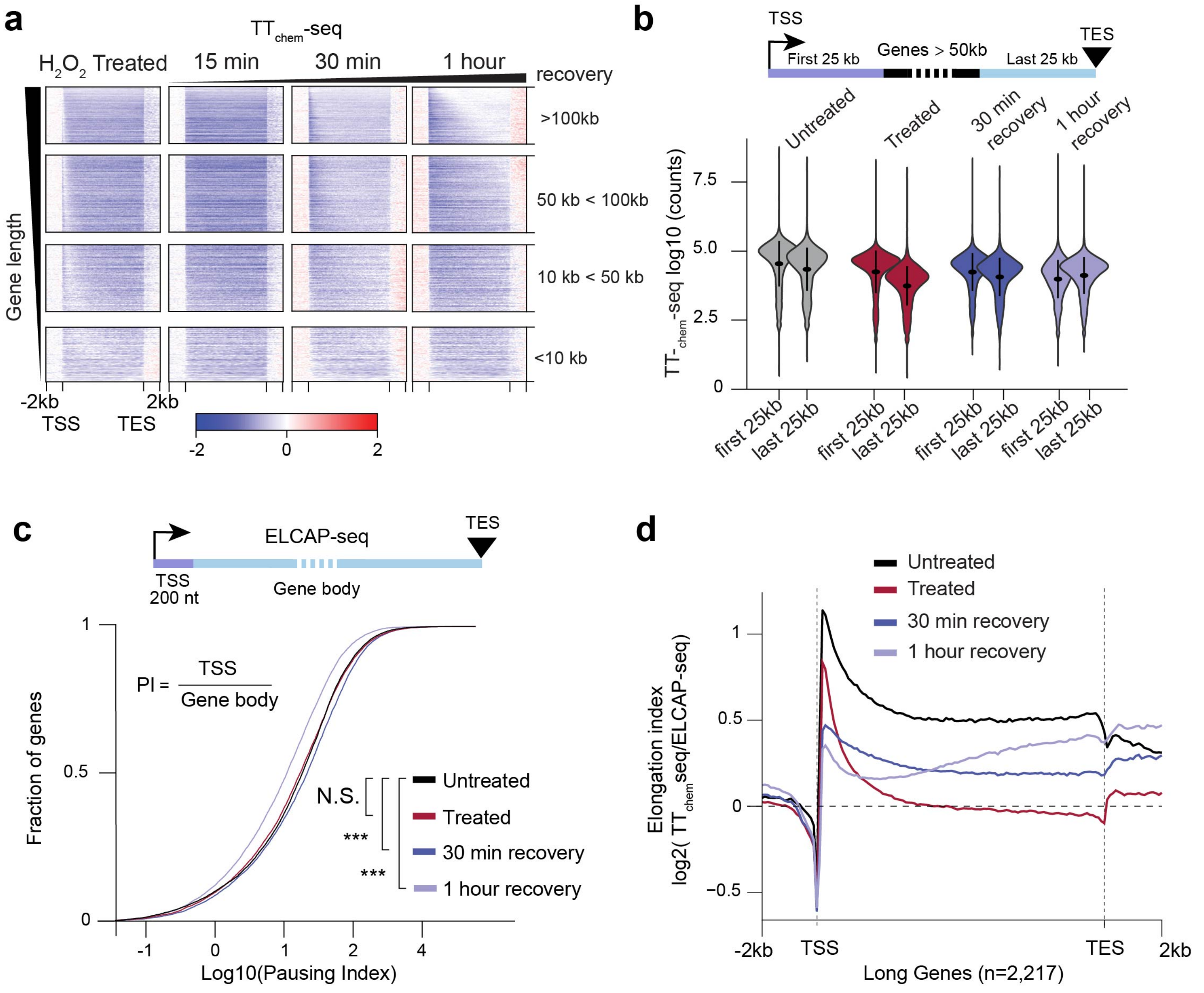


**Supplementary Fig 5**

**Supplementary Fig. 5. Nascent transcription dynamics upon recovery from oxidative stress. a** Heatmaps of $TT_{chem}$-seq signal (normalised to spike-in) in HEK293 cells. The signal is shown as the log2 fold-change for indicated treatment relative to untreated control. Heatmaps are scaled between transcription start site (-2 kb to TSS) and transcript end site (TES +2 kb) Genes are sorted by decreasing sized and separated in four size thresholds. **b** Quantification of $TT_{chem}$-seq signal for the first 25 kb and the last 25 kb of non-overlapping coding genes > 50 kb (n = 5,875). **c** Pausing index (log10 ratio of TSS signal (first 150 nt) relative to remaining gene body signal) of ELCAP-seq signal for all non-overlapping coding genes (n = 17,266). Distribution comparison measured by two-sided Wilcoxon-test, N.S. (p-value > 0.05), *** (p-value < 0.001). **d** Metagene analysis of elongation index, the signal is the log2 fold-change of $TT_{chem}$-seq read counts (normalised to spike-in) signal over RPKM ELCAP-seq signal for respective conditions plotted for long coding genes (> 90 kb, n=2,217).

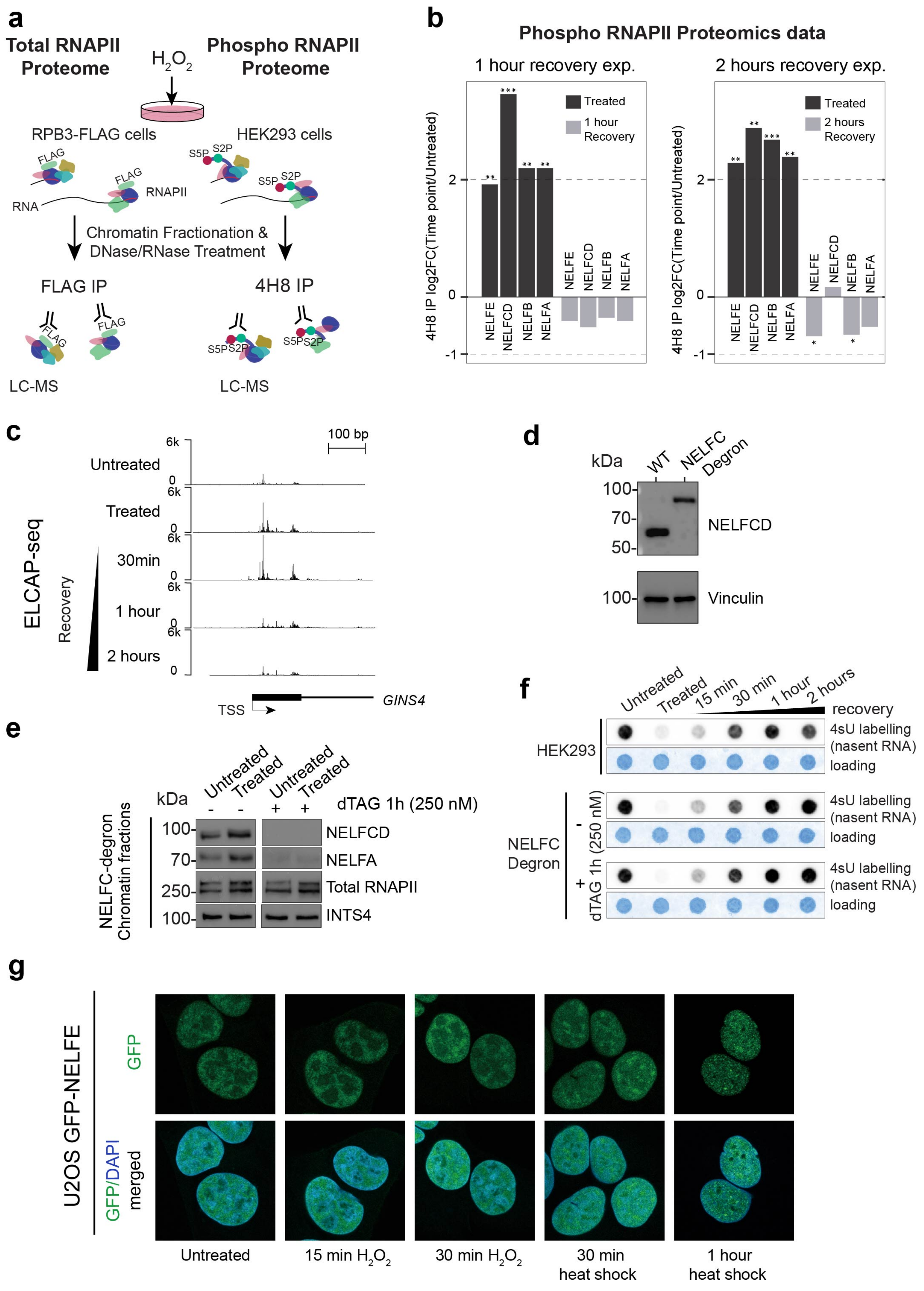


**Supplementary Fig. 6**

**Supplementary Fig. 6. NELF-mediated regulation of RNAPII following oxidative stress. a** Experimental setup used for RNAPII interactome experiments for capture of either total or phosphorylated RNAPII complexes. **b** Log2 fold-change of mass spectrometry average intensities over three technical replicates for individual NELF subunits. Comparison of protein enrichment between treated/1 hour recovery (left) or treated/2 hours recovery (right) over untreated measured by fdr corrected two class unpaired Smar test, * (padj $< 0.05$), ** (padj $< 0.01$),*** (padj $< 0.001$). **c** Single-gene view of ELCAP-seq data for the *GINS4* transcription start site following oxidative stress and recovery. **d** Verification of NELFCD degron cell line. Western blot of WT and NELFCD-degron cells probed with NELFCD antibody and vinculin (loading control). Lower molecular weight is WT NELFC and higher molecular weight band is NELFCD tagged with degron. **e** Western blot of chromatin-enriched cellular fractions from NELFCD-degron cells with or without 1 hour pre-treatment with dTAG-v1 (250 nM) for untreated (no oxidative stress induction) and $H_2O_2$ treated samples. Blots were probed for NELFCD, NELFA, total RNAPII (D8L4Y) and INTS4 as loading control. **f** Nascent transcription levels (4sU dot blot) for HEK293 cells (top) and NELFCD-degron cells (bottom, with and without 1 hour 250 nM dTAG-v1 treatment) treated with 1 mM $H_2O_2$ and indicated times after media replenishment. Staining with methylene blue serves as a loading control. **g** Fluorescence microscopy images of U2OS GFP-NELFE cells showing GFP and DAPI/GFP-merged signals following 15 min $H_2O_2$, 30 min $H_2O_2$ as well as 30 min and 1-hour 43°C heat shock.

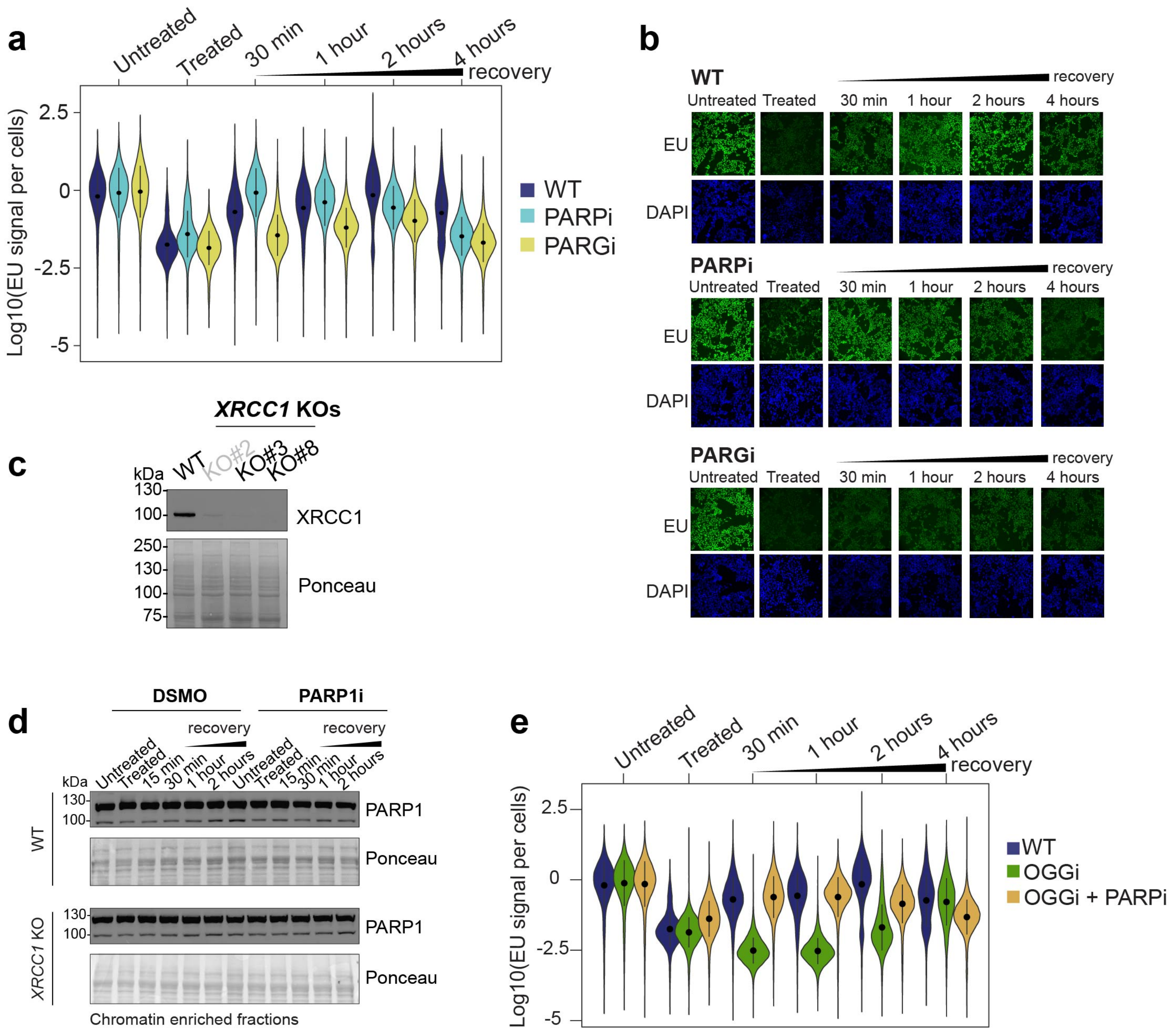


Supplementary Fig. 7

**Supplementary Fig. 7. Role of XRCC1 and OGG1 in RNAPII gene body transcription recovery.** **a** Nascent transcription levels quantified by 5-Ethynyluridine (EU) labelling and single cell quantification by immunofluorescence microscopy (EU-assay). Per cell EU signal is normalised to DAPI and log10 scaled. DMSO (WT), PARPi and PARGi treated HEK293 cells were oxidised-treated with 1 mM $H_2O_2$ and EU-labelled for 30 min. **b** Representative images of global nascent transcription (EU immunofluorescence) from experiments in (a) All image intensities are displayed with same exposure as the untreated control. **c** Western blot validation of HEK293 *XRCC1* KOs. Protein loading is represented by ponceau S staining. The two clones KO#3 and KO#8 were used in parallel for experiment with *XRCC1* KOs. **d** Western blot of PARP1 from chromatin-enriched cellular fractions treated with 1 mM $H_2O_2$ and indicated recovery after media washout in WT HEK293 (top) and *XRCC1* KO cells (bottom). Protein loading is represented by ponceau S staining. **e** Nascent transcription levels quantified by 5-Ethynyluridine (EU) labelling and single cell quantification by immunofluorescence microscopy (EU-assay). Per cell EU signal is normalised to DAPI and log10 scaled. DMSO (WT), OGG1i (same experiment as in Figure 7a) and OGG1i together with PARPi treated HEK293 cells were oxidised-treated with 1 mM $H_2O_2$ and EU-labelled for 30 min.
